# Evolutionary origin, population diversity, and diagnostics for a cryptic hybrid pathogen

**DOI:** 10.1101/2023.07.03.547508

**Authors:** Jacob L. Steenwyk, Sonja Knowles, Rafael W. Bastos, Charu Balamurugan, David Rinker, Matthew E. Mead, Christopher D. Roberts, Huzefa A. Raja, Yuanning Li, Ana Cristina Colabardini, Patrícia Alves de Castro, Thaila Fernanda dos Reis, David Canóvas, Rafael Luperini Sanchez, Katrien Lagrou, Egídio Torrado, Fernando Rodrigues, Nicholas H. Oberlies, Xiaofan Zhou, Gustavo H. Goldman, Antonis Rokas

## Abstract

Cryptic fungal pathogens pose significant identification and disease management challenges due to their morphological resemblance to known pathogenic species while harboring genetic and (often) infection-relevant trait differences. The cryptic fungal pathogen *Aspergillus latus*, an allodiploid hybrid originating from *Aspergillus spinulosporus* and an unknown close relative of *Aspergillus quadrilineatus* within section *Nidulantes*, remains poorly understood. The absence of accurate diagnostics for *A. latus* has led to misidentifications, hindering epidemiological studies and the design of effective treatment plans. We conducted an in-depth investigation of the genomes and phenotypes of 44 globally distributed isolates (41 clinical isolates and three type strains) from *Aspergillus* section *Nidulantes*. We found that 21 clinical isolates were *A. latus*; notably, standard methods of pathogen identification misidentified all *A. latus* isolates. The remaining isolates were identified as *A. spinulosporus* (8), *A. quadrilineatus* (1), or *A. nidulans* (11). Phylogenomic analyses shed light on the origin of *A. latus*, indicating one or two hybridization events gave rise to the species during the Miocene, approximately 15.4 to 8.8 million years ago. Characterizing the *A. latus* pangenome uncovered substantial genetic diversity within gene families and biosynthetic gene clusters. Transcriptomic analysis revealed that both parental genomes are actively expressed in nearly equal proportions and respond to environmental stimuli. Further investigation into infection-relevant chemical and physiological traits, including drug resistance profiles, growth under oxidative stress conditions, and secondary metabolite biosynthesis, highlight distinct phenotypic profiles of the hybrid *A. latus* compared to its parental and closely related species. Leveraging our comprehensive genomic and phenotypic analyses, we propose five genomic and phenotypic markers as diagnostics for *A. latus* species identification. These findings provide valuable insights into the evolutionary origin, genomic outcome, and phenotypic implications of hybridization in a cryptic fungal pathogen, thus enhancing our understanding of the underlying processes contributing to fungal pathogenesis. Furthermore, our study underscores the effectiveness of extensive genomic and phenotypic analyses as a promising approach for developing diagnostics applicable to future investigations of cryptic and emerging pathogens.

## Introduction

There are an estimated 1.7 million deaths per year and greater than 150 million cases of severe fungal infections worldwide ^1^. Beyond the profound toll of lives lost, fungal pathogens pose a substantial economic burden conservatively estimated to be US$11.5 billion but may be as high as US$48 billion ^2–4^. The impact of fungal pathogens to human welfare is also likely to increase due to global climate change ^5^. The importance of fungal diseases was recently acknowledged by the establishment of the first ever priority pathogen list by the World Health Organization ^6^, and it has risen in prominence as a prevalent cause of co-infection of COVID-19 patients ^7,8^.

Even though a great deal is known about several aspects of fungal pathogenicity, we know surprisingly little about the repeated evolution of pathogenicity in fungi and the traits and genetic elements that contributed to it ^9^. Evolutionary approaches have not only proven to be highly informative in understanding how pathogens differ from their non-pathogenic close relatives ^10–13^ but also revealed the presence of cryptic species, defined as species that are genetically distinct from known species but difficult to distinguish using morphology ^14^. Cryptic pathogens have been identified in several fungal genera that contain pathogens, including *Cryptococcus*, *Aspergillus*, *Histoplasma*, and *Fusarium* ^12,15–19^.

Recently, we used short-read genome sequencing data from a sample of six clinical isolates to show that *Aspergillus latus* is a novel cryptic fungal pathogen ^20^. *A. latus* arose via allodiploid hybridization between *A. spinulosporus* and a close, unknown relative of *A. quadrilineatus*, wherein (nearly) the entire genomes of both parents were combined during hybridization. However, the small number of highly fragmented short-read genome assemblies left numerous questions unanswered—such as the timing and number of hybridization events, the prevalence of recombination, and the genetic diversity and phenotypic variation in the species. Moreover, *A. latus* lacks molecular and physical diagnostics to facilitate faithful identification in clinical settings, hindering our understanding of the epidemiology and burden of disease.

To further understand the biology and evolutionary origins of the cryptic fungal pathogen *A. latus*, we generated high-quality genome assemblies using long- and short-read sequencing technologies on a substantially expanded set of 43 globally distributed isolates (40 clinical isolates; three type strains) from section *Nidulantes* of the genus *Aspergillus*. We found that 20 clinical isolates correspond to *A. latus* and the remaining 20 clinical isolates to *A. spinulosporus* (8), *A. quadrilineatus* (1), or *A. nidulans* (11); one previously published strain of *A. latus* was included in the final dataset of 44 isolates ^20^. *A. latus* likely arose from either one or two hybridization events approximately 15.4 to 8.8 million years ago, has a closed pangenome, and exhibits substantial genetic diversity among gene families and biosynthetic gene clusters. Transcriptomics revealed both parental genomes are actively expressed and respond to environmental perturbations. Profiling of each species across infection-relevant chemical and physiological traits suggests hybridization contributes to a unique phenotypic profile. Comprehensive genomic and phenotypic characterization of *A. latus* enabled us to develop five diagnostic markers (three genomic and two phenotypic) that distinguish *A. latus* from closely related species. Characterization of *A. latus*, an understudied cryptic fungal pathogen, expands our understanding of the origins of fungal pathogenicity, establishes a broadly applicable workflow for the development of novel diagnostic markers for pathogens, and is a key step toward preventing and combating disease.

## Results

### Genome sequencing and species identification of 40 clinical isolates

The genomes of three type strains (*A. latus* NRRL 200^T^; *A. spinulosporus* NRRL 2395^T^; *A. quadrilineatus* NRRL 201^T^) and 40 clinical isolates (Supplementary Data 1) were sequenced and assembled using both long-(Oxford Nanopore) and short-read (Illumina) technologies. The resulting genome assemblies are highly contiguous with mean N50 and L50 values of 3.09 ± 1.27 Mbp and 13.24 ± 21.19 scaffolds, respectively (Extended Data Fig. 1), an improvement of ∼10-to-14-fold from previously available genome assemblies ^20^. Species determination via molecular phylogenetic analyses of the taxonomically informative loci of β-tubulin and calmodulin (Extended Data Fig. 2 and 3) placed the clinical isolates into the clades of four distinct species: *A. latus* (20), *A. spinulosporus* (8), *A. quadrilineatus* (1) and *A. nidulans* (11). Notably, species determination using MALDI-TOF mass spectrometry, a standard method for microbe identification in clinical laboratories ^21^, revealed variable accuracy for each species (p < 0.001, Fisher’s exact test; Supplementary Data 2); all examined *A. latus* isolates and six out of nine examined *A. nidulans* isolates were misidentified, whereas all *A. spinulosporus* isolates were accurately identified. This finding suggests that *A. latus* infections are likely underreported. A previously sequenced and assembled clinical isolate of *A. latus*, MO46149 ^20^, was added to the dataset resulting in a total number of 44 total isolates: 41 clinical isolates and three type strains (Extended Data Fig. 4).

**Figure 1.**
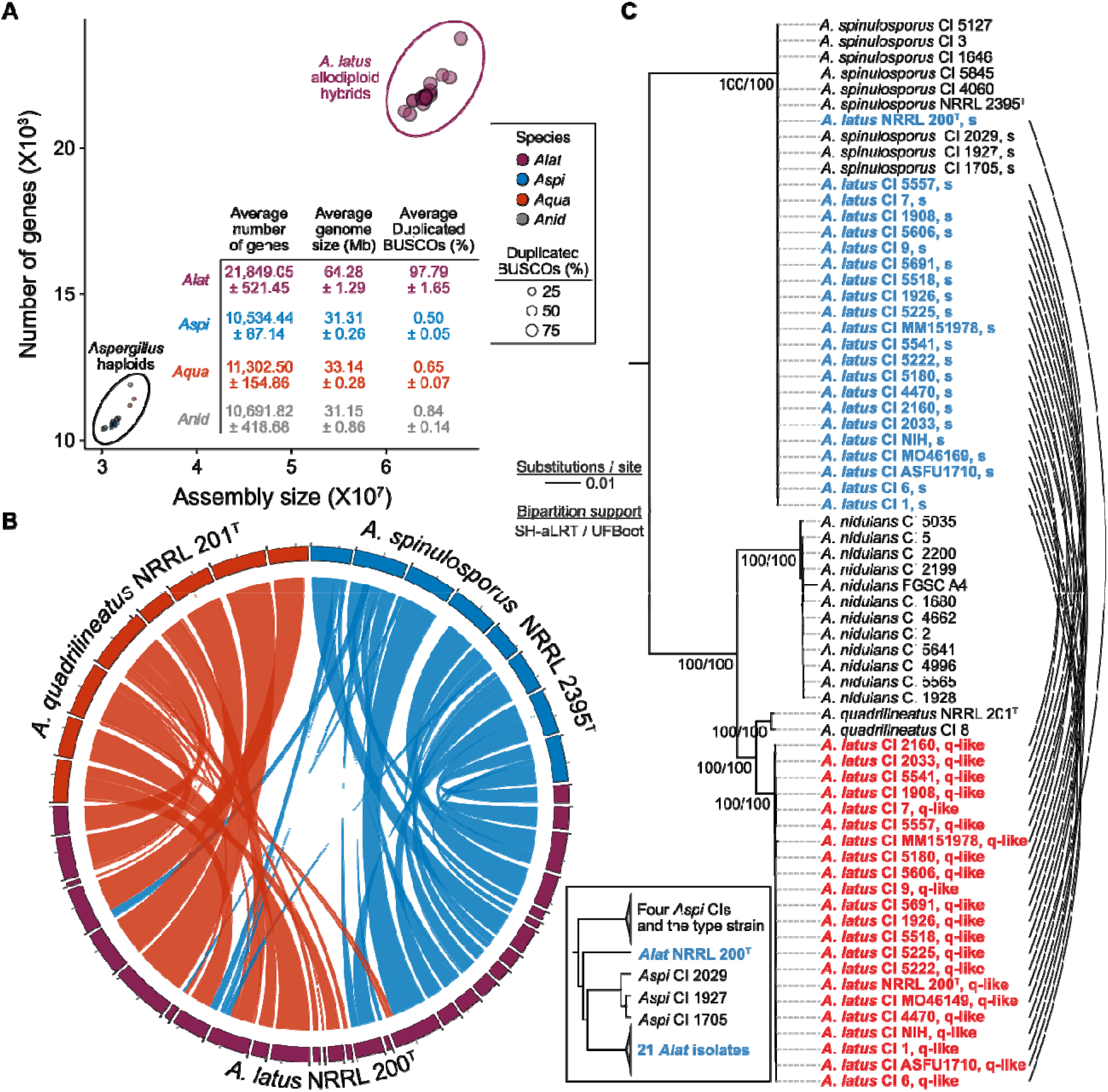
*Aspergillus latus* originated from the hybridization of two closely related species. (A) The genomes of *Aspergillus* hybrids have larger genomes (x-axis), encode more genes (y-axis), and numerous near-universally single copy orthologs (or BUSCO genes) are present in duplicate copies (data point size). The inset table provides averages ± standard deviation for each metric for the diploid genomes of the *A. latus* hybrids (*Alat*; purple) and the haploid genomes of *A. spinulosporus* (*Aspi*; blue), *A. quadrilineatus* (*Aqua*; red), and *A. nidulans* (*Anid*; grey). (B) Synteny analysis between the *A. latus*, *A. quadrilineatus*, and *A. spinulosporus* type strains. Syntenic blocks contain a minimum of 15 genes robustly assigned to one of the parental species. Scaffolds greater than 100 kilobases are depicted. (C) Phylogenomic tree of the 40 clinical isolates and type strains for each species. Note that the genomes of the hybrid isolates and the *A. latus* type strain were split into their two subgenomes (blue: *A. spinulosporus* subgenome; red: *A. quadrilineatus*-like subgenome). Links connect subgenomes from the same isolate. The inset phylogeny highlights the lack of monophyly among *A. latus* isolates *A. spinulosporus* subgenome.

**Figure 2.**
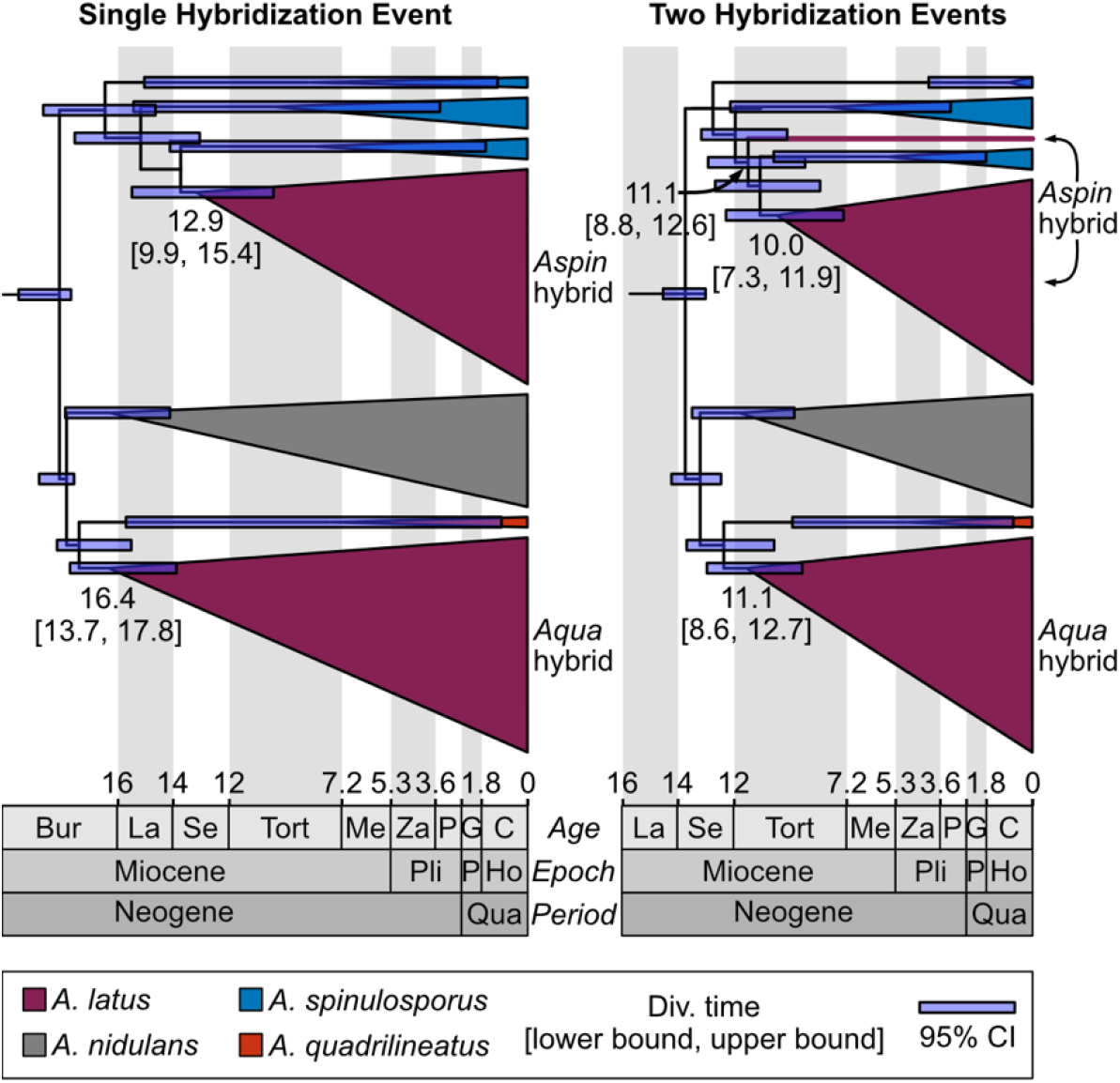
*A. latus* hybrids likely originated approximately 11-13 million years ago. Relaxed molecular clock analyses of the origin of each parental subgenome for the evolutionary scenarios of one- and two-hybridization events (left and right, respectively). Under the hypothesis of a single hybridization event, the *A. spinulosporus* and *A. quadrilineatus*-like subgenomes arose 12.9 and 16.4 million years ago, respectively. Under the evolutionary scenario of two hybridization events, the *A. spinulosporus* and *A. quadrilineatus*-like subgenomes arose 11.1 million years ago, with a second hybridization event 10.0 million years ago. Node bars represent 95% confidence intervals (CI). Lower and upper bound confidence intervals (depicted as: [lower bound, upper bound]) are written for nodes of interest. The phylogeny for the two-hybridization event hypothesis is the same as depicted in Figure 1C; the phylogeny for the single hybridization event hypothesis stems from forcing monophyly of the *A. spinulosporus* subgenomes.

**Figure 3.**
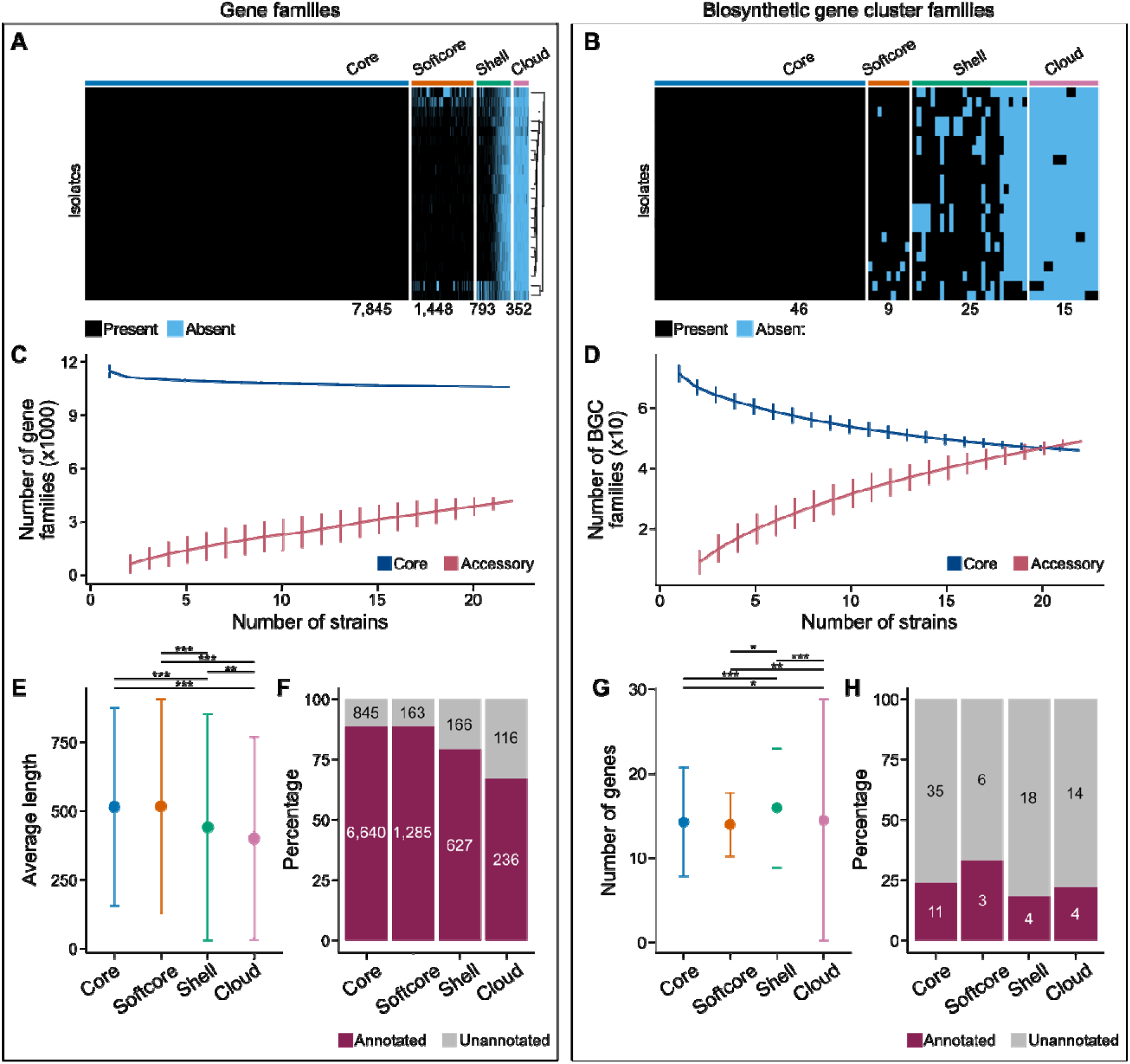
Differing patterns of diversity among gene families and biosynthetic gene clusters in hybrid *A. latus* genomes. (A) Among a total of 10,078 orthologous groups of genes, 7,485 are categorized as core (present in 100% of isolates; N=22) and 2,593 as accessory (present in <100% of isolates; N<22); among these accessory genes, 1,448 are softcore (present in ≥ 95% and <100%; N=21), 793 are shell (5-95% of isolates; 21>N≥2), and 352 are cloud (present in less than 5% of isolates; N=1). (B) Among 95 biosynthetic gene cluster families (BGCFs), 46 are categorized as core and 49 as accessory (9 are softcore, 25 are shell, and 15 are cloud). (C) The number of accessory gene families is increasing as the number of strains increases, suggesting the pangenome is open. (D) The number of accessory BGCFs substantially increases with additional isolates suggesting the pangenome of BGCs is open. Notably, the core genome is larger among gene families, whereas the pangenome is larger among BGCFs. (E) Protein sequence lengths differed among gene categories wherein core and softcore genes are longer than shell and cloud genes. (F) As genes were less frequently observed among isolates, they were also functionally annotated less frequently. (G) The number of genes in BGCs different per category wherein softcore BGCs tend to be smaller than BGCs categorized as core, shell, and cloud. (H) Few BGCs are predicted to make known secondary metabolites.

**Figure 4.**
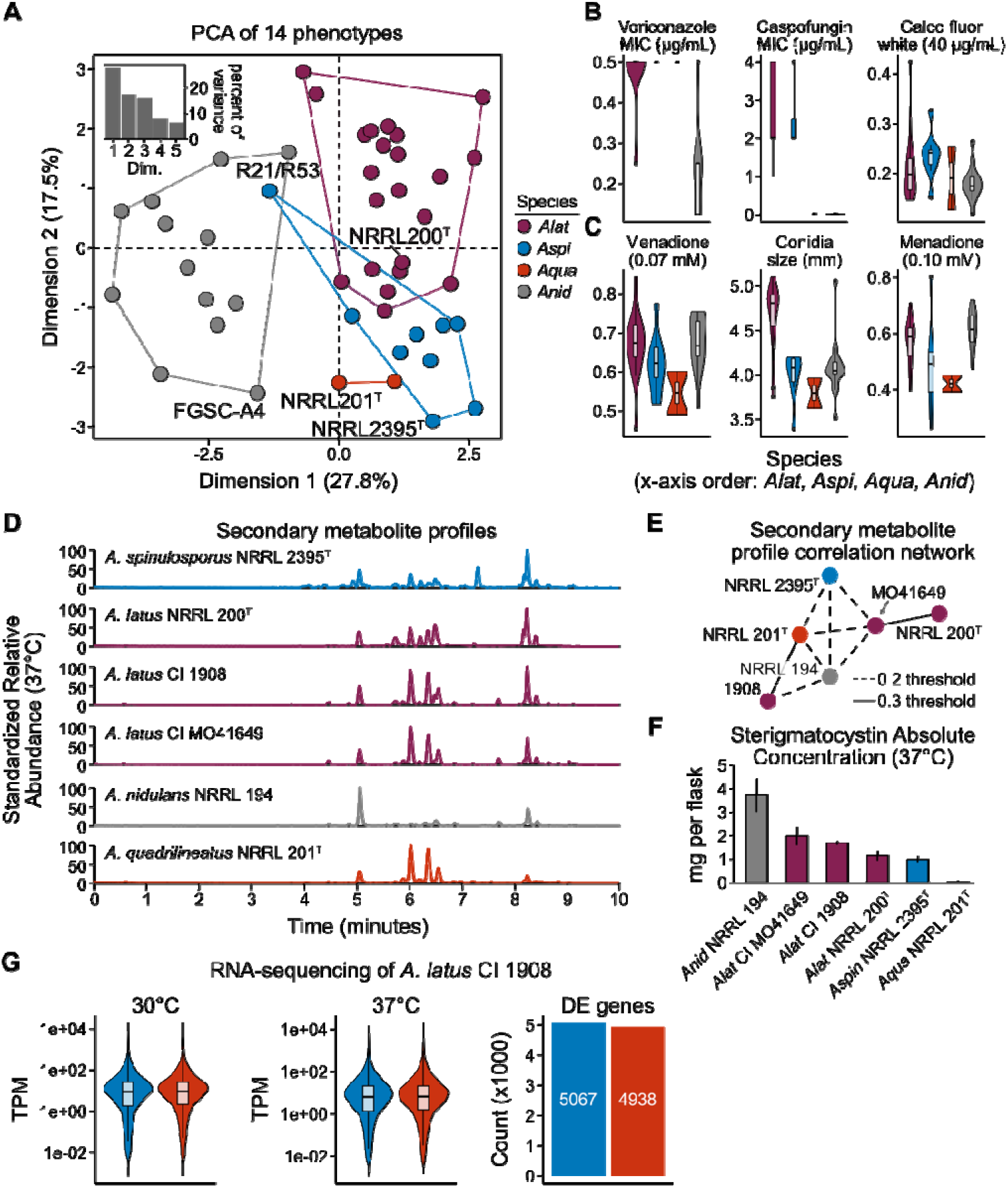
Phenotypic profiling suggests that *A. latus* clinical isolates differ from isolates of closely related species in diverse infection-relevant traits. (A) Principal component analysis of 14 infection-relevant traits reveals each species is phenotypically distinct. Note, some *A. latus* isolates are like some *A. spinulosporus* isolates. (B) Distributions of phenotypic profiles for the three traits that contribute most to the variance along the first and (C) second dimensions. (D) Secondary metabolite profiling of six representative strains and (E) correlation analysis reveals variation between species and within *A. latus*. (F) Absolute concentration of the mycotoxin sterigmatocysin in six strains from four species. (G) Transcriptomics of *A. latus* CI 1908 reveal near-equal RNA abundance between the parental genomes (*A. spinulosporus* in blue; *A. quadrilineatus*-like in red). Moreover, the number of differentially expressed genes between 30°C and 37°C is similar.

### Hybridization gave rise to *A. latus* approximately 12 million years ago

Genome features, phylogenomics, and macrosynteny provide unequivocal support for a hybrid origin of *A. latus*. The genomes of *A. latus* isolates are approximately twice the size, encode twice the number of genes compared to closely related species in section *Nidulantes*, and encode duplicated copies of near-universally single-copy orthologs (or BUSCO genes) suggestive of a diploid genome (Fig. 1A; Extended Data Figs. 1 and 5). Macrosynteny analysis between *A. spinulosporus* NRRL 2395^T^, *A. quadrilineatus* NRRL 201^T^, and *A. latus* NRRL 200^T^ revealed large genomic segments that are syntenic with *A. spinulosporus* and *A. quadrilineatus*, which are indicative of a history of hybridization (Fig. 1B). Further evidence of hybridization was identified using phylogenomics — specifically, the topologies of 3,101 BUSCO genes present in two copies among *A. latus* hybrids showed evidence of one BUSCO gene originating from *A. latus* and the other a close, but unknown, relative of *A. quadrilineatus* (Fig. 1C; Extended Data Fig. 6), consistent with previous findings ^20^.

**Figure 5.**
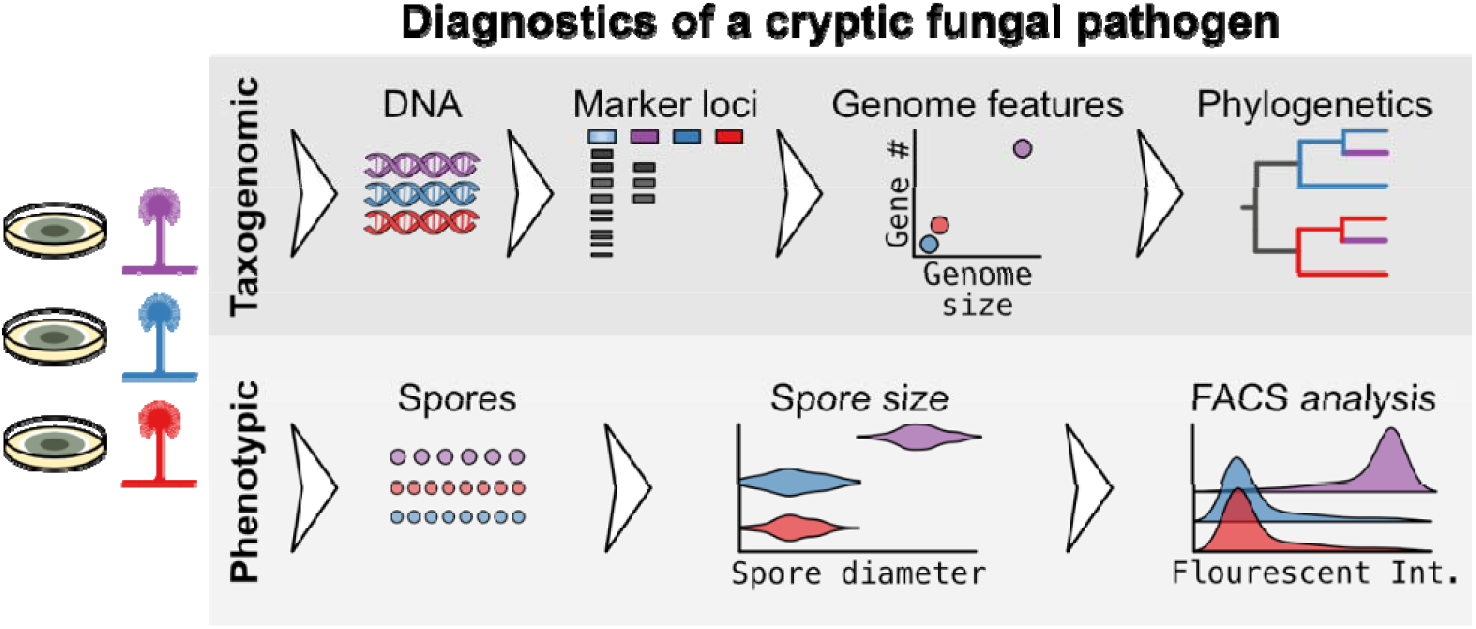
Proposed diagnostics for reliable identification of *A. latus*. *A. latus* (purple), *A. spinulosporus* (blue), and *A. quadrilineatus* (red) are indistinguishable in culture, and we provide evidence of lack of sufficient accuracy for MALDI-TOF-based species identification. Genomic and phenotypic analyses revealed five markers that can be used for diagnostics, each requiring different levels of analytical complexity, thereby accommodating variation in resources available in clinical settings. Diagnostic markers can be divided into two categories. Taxogenomic markers include three gene families uniquely present in *A. latus* isolates, two genome features (i.e., gene number and genome size), and phylogenetic/phylogenomic methods. A prerequisite for the use of these markers is genome sequencing. Phenotypic markers include spore size and genome size (estimated using Fluorescence-Activated Cell Sorting or FACS), which require isolating asexual spores.

The denser sampling of isolates allowed us to investigate the number and timing of hybridization events that gave rise to *A. latus*. The maximum likelihood phylogeny of our phylogenomic dataset, which includes all the parental subgenomes in the *A. latus* isolates as well as all *A. spinulosporus*, *A. quadrilineatus*, and *A. nidulans* isolates, suggests that *A. latus* originated via two distinct hybridization events – one for the *A. latus* NRRL 200^T^ and one for all other *A. latus* isolates (Fig. 2). However, there is low support for the divergence of *A. latus* NRRL 200^T^ from other isolates (SH-aLRT = 78.9; UFBoot = 85; Fig. 1C; Extended Data Fig. 7). Analyses of a subsample of the total phylogenomic data matrix (N_genes_ = 3,101) filtered to include only those genes with high taxon occupancy (both parental sequences present for at least 19 hybrid isolates; N_genes_ = 1,183) also weakly supported the occurrence of two hybridization events (SH-aLRT = 93.7; UFBoot = 80; Extended Data Fig. 7). An approximately unbiased test was used to test whether a constraint tree where all the *A. spinulosporus* subgenomes of *A. latus* isolates were monophyletic had a significantly lower likelihood score than the unconstrained maximum likelihood tree. The likelihood scores of the two trees were not significantly different in both data matrices (ΔL_complete data matrix_ = 171.47, p = 0.071; ΔL_subsampled data matrix_ = 157.98, p = 0.060). These results were consistent using different methods of topology testing (Supplementary Tables 1 and 2). Therefore, we cannot rule out a single hybridization event.

Relaxed molecular clock analyses suggest *A. latus* arose approximately 13-11 million years ago during the Miocene (Fig. 2). Divergence time estimates differed for the evolutionary scenario of one or two hybridization events. Specifically, for the two-hybridization events hypothesis, the *A. spinulosporus* and *A. quadrilineatus*-like subgenomes arose 11.1 (12.6 – 8.8) million years ago and a second hybridization event occurred 10 (11.9 – 7.3) million years ago; for a single hybridization event hypothesis, the *A. spinulosporus* and *A. quadrilineatus*-like subgenomes arose 12.9 (15.4 – 9.9) million years ago and 16.4 (17.8 – 13.7) million years ago, respectively.

### Parental genomes rarely recombine

The new genome assemblies and the high nucleotide sequence divergence (7%) of the subgenomes enabled us to examine whether recombination occurs between the subgenomes of *A. latus*. Using the *A. latus* NRRL200^T^ genome as a reference, we found that that its two subgenomes display little evidence of recombination, as measured by scaffolds containing syntenic blocks of 15 or more consecutive genes all assigned to one or the other parent (Fig. 1B). Specifically, we found that 88.30% (1,057 / 1,197) of scaffolds did not show signatures of recombination between the parental subgenomes (Supplementary Data 3. Among the 11.70% (140 / 1,197) scaffolds that did show evidence of recombination, 118 had two syntenic blocks, one from each parent; 18 had three syntenic blocks; three had four syntenic blocks, zero had five syntenic blocks; and one had six syntenic blocks (Extended Data Fig. 8; Supplementary Data 4).

Examination of the relationship between the frequency of recombination and the genome assembly quality of the *A. latus* isolates revealed that the number of scaffolds with evidence of recombination is significantly correlated to the number of scaffolds in a genome assembly (ρ = 0.57, p = 0.007, Spearman rank correlation) and the genome assembly N50 (ρ = -0.48, p = 0.027, Spearman rank correlation) (Extended Data Fig. 9). Since the number of scaffolds with evidence of recombination increased as genome quality decreased, at least some of the observed recombination is likely an artifact stemming from genome assembly errors, suggesting that the *A. latus* hybrid genomes display little, if any, evidence of recombination between the parental subgenomes.

### The pangenome of *A. latus* is large and that of biosynthetic gene clusters even larger

To examine the pangenome of *A. latus*, we assigned orthologous groups of genes (a proxy for gene families) to four different bins of occupancy: core (present in all *A. latus* isolates; N=22) and accessory (present in <100% of *A. latus* isolates); accessory genes were assigned as softcore (present in nearly all isolates; N=21), shell (present in 5-95% of isolates; 2≤N<21), or cloud (present in less than 5% of isolates; N=1). Across 10,078 gene families, 7,485 (74.27%), 1,448 (14.37%), 793 (7.87%), and 352 (3.49%) were determined to be core, softcore, shell, and cloud, respectively (Fig. 3A). We then used the same approach to define the pangenome of biosynthetic gene clusters (BGCs) involved in secondary metabolism (hereafter referred to as the BGCome). Assigning BGCs to BGC families revealed that 46 (48.42%), 9 (9.47%), 25 (26.32%), and 15 (15.79%) BGC families were assigned to the core, softcore, shell, and cloud, respectively (Fig. 3B).

Examination of the relationship between the number of gene families or BGCs in the core and accessory (softcore, shell, and cloud) pangenome underscored their differing evolutionary dynamics. Core genes are substantially more common than accessory genes in the *A. latus* pangenome (Fig. 3C). In contrast, the accessory BGCome is larger than the core (Fig. 3D). Although both the pangenome and the BGCome remain open, these findings suggest that the evolutionary trajectory of the BGCome is distinct from that for the entire pangenome. Further sampling will enable more precise estimates of the size of the core and accessory parts of the *A. latus* genome.

Proteins encoded by the core and softcore gene families differ in protein sequence lengths (χ2 = 165.57, p < 0.001, df = 3; Kruskal-Wallis Rank Sum test) (Fig. 3E). Specifically, the average protein sequence length among core and softcore genes is 515.25 ± 369.06 and 517.45 ± 389.26 amino acids, respectively, and are significantly longer than shell and cloud genes that have an average length of 441.94 ± 412.28 and 401.68 ± 369.06 amino acids, respectively (p < 0.001 for both comparisons; Dunn’s test, Benjamini-Hochberg (BH) multi-test correction). Shell genes are also longer than cloud genes (p = 0.004; Dunn’s test, BH multi-test correction). Core and accessory genes also differed in the proportion of genes that could be functionally annotated, wherein a greater proportion of genes were annotated in the core and softcore gene families and fewer in the shell and cloud (Fig. 3F).

Core and accessory BGCs significantly differed in the number of genes encoded in each (χ2 = 166.19, p < 0.001, df = 3; Kruskal-Wallis Rank Sum test) (Fig. 3G). Specifically, softcore BGCs, which have an average of 10.69 ± 3.99 genes, encode significantly fewer genes than core, shell, and cloud BGCs, which have an average of 14.45 ± 4.46, 15.04 ± 6.71, and 15.10 ± 7.04 genes, respectively (p < 0.001 for all comparisons; Dunn’s test with BH multiple test correction). The products, if any, of most BGCs are not known, suggesting *A. latus* isolates may be capable of producing novel secondary metabolites; alternatively, numerous BGCs encoded in the *A. latus* genomes may be nonfunctional (Fig. 3H).

### Phenotypic profiles of hybrid pathogen isolates are distinct

Hybridization can change the phenotypic profile of organisms ^22^. Extensive phenotyping of infection-relevant traits (i.e., growth under cell wall perturbation and integrity stresses, growth under oxidative stress, growth at 44°C, 37°C, and 30°C, susceptibility to voriconazole, amphotericin B, and caspofungin, growth in minimal media, asexual spore size, and production of cleistothecia) across all isolates including a synthetic diploid of *A. nidulans*, strain R153XR21, revealed that hybrids were distinct from the other three species, but were most similar to *A. spinulosporus* (Fig. 4A). Drug resistance and a cell wall stressor were the largest contributors to the variance along the first dimension in principal component space (Fig. 4B; Extended Data Fig. 11), whereas oxidative stress and asexual spore size contributed most to the variance along the second dimension (Fig. 4C). Notably, 14 *A. latus* and five *A. spinulosporus* isolates had high drug resistance to caspofungin compared to reference isolates of *A. nidulans* (varying from 2.0 to > 4.0 ug/mL and 0.03 ug/mL, respectively). Variation among the species was also observed in an invertebrate moth model of disease (p = 0.04; Log-Rank test) (Extended Data Fig. 12; Supplementary Data 6). Variation among clinically-relevant traits suggests that each species may warrant unique strategies to prevent and combat disease.

Examination of secondary metabolite profiles among six isolates—three *A. latus* isolates and single representatives from *A. spinulosporus*, *A. quadrilineatus*, and *A. nidulans*—revealed that *A. latus* hybrids were qualitatively more similar to *A. quadrilineatus* (Fig. 4D). Correlation analysis revealed that *A. latus* CI 1908 was more similar to *A. quadrilineatus* NRRL 201^T^ than to other strains (Fig. 4E). Examination of the absolute concentration of a mycotoxin, sterigmatocystin, in the same panel of six isolates revealed that *A. latus* hybrids produced more toxin than either parental species (*A. spinulosporus* and *A. quadrilineatus*) but less than *A. nidulans* (Fig. 4F). Together with other phenotyping, these results suggest that *A. latus* hybrids are phenotypically distinct from other species but are more like one parent for some traits, more like the other parent for others, and differ from both parents in yet others.

We next determined if *A. latus* gene expression patterns were biased toward one or the other parent. RNA-sequencing of *A. latus* CI 1908 at 30°C and 37°C revealed nearly equal transcript abundances in both parental genomes (Fig. 4G). For example, at 37°C the *A. spinulosporus* and *A. quadrilineatus*-like subgenomes had average transcript per million (TPM) values of 47.72 ± 355.86 and 49.44 ± 369.27, respectively; a similar observation was made for transcript abundances at 30°C. Differential expression analysis between the two conditions revealed similar numbers of differentially expressed genes in each parental genome (5,067 and 4,938 in the *A. spinulosporus* and *A. quadrilineatus* subgenomes, respectively; p < 0.01 and fold change > 2). Examining the estimated codon usage bias of each parental genome (Extended Data Fig. 13) corroborates this finding, revealing that both subgenomes are nearly equally optimized. These findings suggest that parental genomes contribute to organismal function and respond to environmental perturbations.

### Novel diagnostics for reliable *A. latus* identification

Since all *A. latus* isolates in the present study were misidentified by traditional methods (Supplementary Data 2), we used the newly generated genomes to identify *A. latus*-specific molecular markers and traits for potential clinical use (Fig. 5). Although best used in combination, each marker is designed to be sufficient to differentiate these species. Notably, markers vary in complexity (owing to anticipated variation in accuracy and precision), enabling clinicians with limited access to resources (ranging from microscopes to high-performance computing clusters) to identify *A. latus*.

Among taxogenomic markers, we propose three lines of evidence that can be used to help diagnose *A. latus* infections in a clinical setting: 1) three marker loci present in all *A. latus* strains but absent in *A. spinulosporus*, *A. quadrilineatus*, and *A. nidulans* (Supplementary Data 7); polymerase chain reactions for these loci can help identify *A. latus* isolates, 2) whole-genome sequencing; *A. latus* isolates have larger genome sizes and gene repertoires than other *Aspergillus* species (Fig. 1A), and 3) hybridization among single-locus trees, including taxonomically informative loci (Fig. 1C).

Among phenotypic markers, we propose two lines of evidence that point toward an *A. latus* infection, which can be discerned from spores. Specifically, *A. latus* spores are larger (Fig. 4C) resulting from larger genome sizes ^20^. Continuing to leverage genome size variation, Fluorescence-Activated Cell Sorting (or FACS) analysis will uncover more DNA content in the *A. latus* spores than in other *Aspergillus* species (Supplementary Data 8). These taxogenomic and phenotypic markers provide the first diagnostics for *A. latus* infections, which may help elucidate the epidemiology, and clinical burden of this cryptic pathogen, a key step in combating and preventing fungal infections.

## Discussion

Our study generated and analyzed extensive amounts of genomic, transcriptomic, chemical, and phenotypic data to shed light on a cryptic fungal pathogen’s evolutionary history, genetic diversity, and the signature of hybridization across infection-relevant traits. Evolutionary analyses indicate that *A. latus* arose approximately 12 million years ago via one or two hybridization events (Figs. 1 and 2). If *A. latus* originated via two events, our analyses suggest that both must have occurred around the same time.

Examination of the *A. latus* pangenome revealed that it harbors considerable genetic diversity, with 74.27% (7,485 / 10,078) of the pan-genes being core (Fig. 3). For comparison, the size of the *Candida albicans* pangenome is 5,919 - 6,054 gene families and 5,432 (89.73 - 91.77%) of its genes are core (present in all isolates) ^23^, whereas the *A. fumigatus* pangenome is more open and only 69.34% (7,563 / 10,907) of pan-genes are core ^24,25^. In contrast to pan-genes, the accessory BGCs outnumber core BGCs — 51.58% of BGCs are accessory — corroborating previous reports that biosynthetic gene clusters evolve rapidly ^13,26^. The *A. latus* BGCome is less diverse than the *A. fumigatus* BGCome —specifically, 46 (48.42%) of *A. latus* BGCs are core, whereas 30.55% (11 / 36) of *A. fumigatus* BGCs are core ^27^. A substantial portion of pan-genes, especially in the BGCome, remain uncharacterized and may represent an untapped source of novel biology and chemistry, especially in the context of an allodiploid organism.

*A. latus* has a distinct phenotypic profile compared to parental species and *A. nidulans*, which is one of the species that *A. latus* has been previously mistyped as ^20^. We hypothesize that some phenotypes may be additive—for example, *A. latus* produces the mycotoxin sterigmatocystin in an amount roughly equal to the sum of the amounts produced by its parental species (Fig. 4F). In other cases, the hybrids are more like one parent over the other. For example, across all infection-relevant traits examined, *A. latus* is closer to *A. spinulosporus* than to *A. quadrilineatus*; however, this observation may partly be explained by the fact that *A. spinulosporus* is one of the parental species whereas *A. quadrilineatus* is a close relative of the other parental species (and not the actual parental species). In contrast, *A. latus* CI 1908 has a secondary metabolite profile more like *A. quadrilineatus* than *A. spinulosporus*. Similarity to one parent over the other may be partly due to dominant genetic effects. These observations corroborate previous findings that complex additive and non-additive effects contribute to hybrid phenotypic profiles in fungi ^28^ and plants ^29^.

The expression patterns of *A. latus* subgenomes differ from those previously observed in other allodiploid hybrids. Whereas both *A. latus* subgenomes are actively transcribed and nearly equally respond to environmental perturbations (Fig. 4G), the plant pathogen *Verticillium longisporum* displays subgenome-specific patterns of gene expression ^30^. Moreover, we observe that *A. latus* genomes tend to maintain allodiploidy whereas *V. longisporum* undergoes haploidization, a common genomic event in the aftermath of hybridization ^30^. Maintaining genetic loci after hybridization may result from selection, as previously proposed for *Coccidioides* fungi ^31^. Whether selection is acting to maintain considerable portions of both parental genomes in *A. latus* hybrids remains an open question.

Accurate identification of pathogenic microbes can help inform disease management strategies. Our extensive population-level characterization of genomic and phenotypic diversity in *A. latus* facilitated the development of the first set of diagnostics for this cryptic pathogen. Importantly, these markers were designed to accommodate for variation in resource availability, facilitating their use in broad clinical settings across the globe. Markers can be grouped into two broad categories: taxogenomic and phenotypic. Taxogenomic markers include gene families uniquely present in *A. latus*, genome features, and gene families present in two (divergent) copies in *A. latus* (Figs. 1 and 5; Supplementary Data 7). Phenotypic markers include spore size (which is positively correlated with genome size) and FACS analysis of spores (Figs. 4 and 5; Supplementary Data 7). These markers can reliably distinguish *A. latus* from other species in section *Nidulantes* and hold promise for illuminating the epidemiology and clinical burden of *A. latus* paving the way for better informed disease management strategies.

This study expands our understanding of the evolutionary origins and phenotypic impact of hybridization in an enigmatic cryptic fungal pathogen of humans via extensive genomic and phenotypic analysis. Moreover, these analyses facilitated the development of the first set of diagnostic markers for the reliable identification of *A. latus* pathogens. Population sequencing and phenotypic analyses emerge as a promising approach for developing diagnostics, which may help rapidly develop markers for future cryptic and emerging pathogens.

## Materials and Methods

### Genomic DNA extraction

The high molecular weight DNA was extracted as follows. Fungal conidia were inoculated in liquid media and let to grow at 37°C for 16h under agitation. Further, the mycelia were filtered through miracloth, freeze dried, and disrupted by grinding in liquid nitrogen. The ground material was transferred to a 50 mL Falcon tube, lysed in 10mL of lysis buffer (3.75 ml Buffer A [3.19 g Sorbitol, 5 mL 1M Tris-HCl (pH 9), 0.5 mL 0.5M EDTA (pH 8) and ddH2O up to 50 mL], 3.75 ml Buffer B [10 mL 1M Tris-HCl (pH 9), 5 mL 0.5M EDTA, 5.84 g NaCl, 1 g CTAB and ddH2O up to 50 mL], 1.5 mL 5 % Sarkosyl; 1 mL 1 % PVP; 100 μl Proteinase K) and incubated 30 min at 65°C. Subsequently, 3.35mL of 5 M Potassium acetate (pH 7.5) were added and the samples were incubated on ice for 30 min. After 30 min centrifugation at 5,000 g and 4°C, the supernatants were transferred to a new falcon tube. Subsequently, the samples were treated with Phenol:Chloroform:Isoamylalcohol (25:24:1), tubes were centrifuged, the aqueous phase (∼8mL) was transferred to a new tube and treated with 100 μL RNase A (10 mg/ml) for 60 min at 37°C. The genomic DNA was precipitated by the addition of 1/10 volume of 3M Sodium acetate and 1 volume ice-cold 96% Ethanol and centrifuged 30 min at 10,000 g, 4°C. Finally, the pellet was finally washed with 70% ethanol, dried, resuspended in 500 μL TE and stored at -20°C until further use. The DNA concentration was estimated using a Qubit fluorometer. All other Aspergillus genomic DNA extraction were performed according to Goldman and colleagues 1995 ^32^. The resulting DNA was sequenced using Oxford Nanopore and Illumina technologies at Nextomics Biosciences in Wuhan, China. This resulted in 10,565,680 Nanopore reads (152,924,029,920 total bases) with an average length of 15,492.31 ± 4,991.34 base pairs and 1,145,484,780 Illumina reads with a length of 150 base pairs (171,822,717,000 bases).

### Genome assembly and annotation

Each isolate was assembled using two different approaches. First, a long sequencing read-based assembly was generated by NextDenovo, v2.3.0 ^33^, using Oxford Nanopore sequencing data alone. The assembly was then polished by Nextpolish, v1.3.0 ^34^, using both long- and short-sequencing read data. Second, a hybrid assembly was generated by MaSuRCA, v3.4.1 ^35^, using both long- and short-sequencing data. Prior to assembly, reads were quality-trimmed using fastp, v0.12.6 ^36^, with default settings. The two assemblies were compared for their assembly size, scaffold number, and scaffold N50 value. For the *A. latus* isolates, the hybrid assemblies were selected as they were about twice in size as the corresponding Nanopore-based assemblies. For other isolates, the Nanopore-based assemblies were selected as they were better in continuity. For genome annotation, all genome assemblies were first softmasked for repetitive sequences using RepeatMasker, v4.1.0 (http://www.repeatmasker.org), (“-species” option set to “Aspergillus”). Gene models were generated using BRAKER, v2.1.4 ^37^, which combines ab-initio gene predictors (AUGUSTUS, v3.3.3 ^38^, and GeneMark-ES, v4.59 ^39^) and homology evidence (all Eurotiomycetes protein sequences in the OrthoDB, v10 ^40^, database).

To examine the quality of all genome assemblies examined, diverse metrics that describe genome assembly and gene content completeness were determined. Genome assembly metrics—such as assembly size, the number of scaffolds, N50, L50, and others—were calculated using BioKIT, v0.0.9 ^41^. Gene content completeness was evaluated using BUSCO, v4.0.4 ^42^ with the Eurotiales dataset (creation date: 2020-08-05) of 4,191 near-universally single copy orthologs (commonly referred to as BUSCO genes) from OrthoDB, v10 ^40^.

### Assigning genes to parent-of-origin

Determining gene parent-of-origin was done using two approaches. For BUSCO genes, a quartet decomposition approach was implemented wherein single-gene phylogenies were pruned to quartets (phylogenies with four leaves) with the type strains of *A. spinulosporus* NRRL2395^T^, *A. quadrilineatus* NRRL201^T^, *A. nidulans* A4, and a single-gene from a newly sequenced isolate (Extended Data Fig. 10). The parent-of-origin was determined to be the species whose sequence exhibited the shortest nodal distance to the sequence of the newly sequenced isolate in the quartet. As expected, genes from haploid isolates determined to be *A. spinulosporus*, *A. quadrilineatus*, and *A. nidulans* using molecular phylogenetics (Extended Data Fig. 2 and 3) were correctly assigned to their respective species (Extended Data Fig. 6) demonstrating the efficacy of this approach. Single-gene phylogenies were constructed by aligning protein sequences of BUSCO genes using MAFFT, v7.402 ^43^, with the “auto” parameter. Nucleotide sequences were threaded onto the protein alignments using the “thread_dna” function in PhyKIT, v1.10.0 ^44^, to create codon-based alignments. The resulting alignments were trimmed with ClipKIT, v1.3.0 ^45^, using the “smart-gap” mode. Single-gene phylogenies were inferred using IQ-TREE 2, v2.0.6 ^19^. The best fitting substitution model was automatically determined according to Bayesian Information Criterion using ModelFinder ^46^. Nodal distances were calculated using the “tip_to_tip_node_distance” function in PhyKIT, v1.10.0 ^44^. This analysis was done for 3,738 BUSCO genes. For all other genes, a reciprocal best blast hit approach was used between the coding sequences of the query hybrid genome and the concatenated coding sequences of *A. spinulosporus* NRRL2395^T^ and *A. quadrilineatus* NRRL201^T^.

### Reconstructing species- and strain-level evolutionary histories

To determine the number of hybridization events and to place the evolutionary history of the hybrid subgenomes in geologic time, a phylogenomic approach was employed. First, 3,101 BUSCO genes with at least 45 sequences out of 72 haploid and diploid subgenomes (62.5% taxon occupancy) were identified. The trimmed codon-based alignments generated during parent-of-origin determination were concatenated together. For hybrid genomes, only sequences from the same subgenome were concatenated together. For example, *A. latus* NRRL200^T^ was represented by two concatenated sequences: one representing the *A. spinulosporus* subgenome and the other representing the *A. quadrilineatus*-like subgenome. The resulting alignment had 5,825,649 sites (2,107,046 parsimony informative sites; 3,229,451 constant sites). Alignment summary statistics were calculated using BioKIT, v0.0.4^41^. The concatenated matrix was used to reconstruct the evolutionary history of all species and isolates using IQ-TREE 2, v2.0.6 ^19^. During tree search, the number of trees in the candidate set was increased from five to ten. The best fitting model was determined using Bayesian Information Criterion values as described above. Bipartition support was measured using two metrics: 5,000 ultrafast bootstrap approximations ^47^ and Shimodaira-Hasegawa-like approximate likelihood ratio test using 1,000 replicates^48^.

To ensure phylogenetic inferences were not sensitive to missing data, a known source of error ^49^, a phylogenomic tree was also inferred using a data matrix constructed from genes with high taxon occupancy values. Specifically, 1,183 BUSCO genes with at least 38 diploid sequences (corresponding to 86.36% occupancy and representation from 19 / 22 diploid isolates) were identified. The resulting sequences were concatenated into a single matrix with 2,228,409 sites (812,508 parsimony informative sites; 1,227,455 constant sites). The same tree search parameters and support metrics (ten candidate trees; 5,000 ultrafast bootstrap approximations; Shimodaira-Hasegawa-like approximate likelihood ratio test with 1,000 replicates) were used as previously described.

### Determining the number of hybridization events

The *A. spinulosporus* parental subgenomes of hybrid isolates was placed into two distinct clades in our phylogenomic analysis (Fig. 1C), suggestive of two hybridization events. To evaluate if a single hybridization event also accurately describes the evolutionary history of *A. latus* hybrids, a topology test was conducted. The log-likelihoods of the inferred phylogeny in Fig. 1C and a maximum likelihood phylogeny where the *A. spinulosporus* subgenomes were constrained to be monophyletic were compared using an approximately unbiased test ^50^ with 5,000 replicates ^51^. The test was done for both the 3,101-gene and the 1,183-gene data matrices. To ensure our results were not sensitive to the testing approach, six other topology tests were also conducted: bootstrap proportions using resampling estimated log-likelihoods ^51^; a one-sided Kishino-Hasegawa and weighted Kishino-Hasegawa test ^52^; a Shimodaira-Hasegawa and weighted Shimodaira-Hasegawa test ^53^; and an expected likelihood weight test ^54^.

### Timetree analysis

To place the evolutionary origins of *A. latus* in geologic time, timetree analysis was conducted using the Bayesian framework implemented in MCMCTree from PAML, v4.9d ^55^ and the 1,183-gene data matrix. Sequences in the 1,183-gene data matrix were first subsampled to those belonging to the section *Nidulantes* (N = 70) and the sister section, *Versicolores* (N = 2). Next, the substitution rate was estimated using a strict clock model (clock = 1) and a general time reversible model with unequal rates and unequal base frequencies ^56^; rate heterogeneity across sites was accounted using a discrete Gamma model with four categories ^57^ (model = 7); the root was point calibrated to 27.55 million years ago—the split between sections *Nidulantes* and *Versicolores* based on previous whole-genome estimates ^58^. The substitution rate was estimated to be 0.07 nucleotide substitutions per site per 10 million years.

Next, the likelihood of the alignment was approximated using a gradient and Hessian matrix—a matrix that describes the curvature of the log-likelihood surface. Estimating a gradient and Hessian matrix requires time constraints, which were set as follows: the origin of *Nidulantes* was constrained to an upper bound of 5.44 million years ago and a lower bound of 16.57 million years ago; the split between *Nidulantes* and *Versicolores* was set to an upper bound of 19.79 million years ago and a lower bound 35.40 million years ago). Time constraints were based on the same study used to point calibrate the root during substitution rate estimation ^58^. Lastly, the resulting gradient and Hessian matrix divergence times were estimated using a relaxed molecular clock (clock = 2). The gamma distribution shape [*a* = (*s* / *s*)^2^] and scale [*b* = *s* / *s* ^2^] are defined by the substitution rate (*s*) and were set to 1 and 13.99, respectively (Supplementary Data 5). Rate variation across branches was accounted for by setting the σ^2^ parameter to “1 4.5.” During Markov Chain Monte Carlo analysis, a total of 5.1 million iterations were run—the first 100,000 observations were discarded and a total of 10,000 samples were obtained by sampling every 500 iterations. The total number of iterations run is 510 times greater than the recommended minimum ^59^. This analysis was conducted for tree topologies representing one- and two-hybridization events.

### Pan-genome analysis

Protein sequences in the dataset were clustered into orthologous groups of genes (a proxy for gene families) using OrthoFinder, v2.3.8^60^ using an inflation value for Markov clustering of 1.5. Sequence similarity searches were done using DIAMOND, v2.0.13.151^61^. The resulting gene families were then binned into categories reflecting gene family occupancy following a previously established protocol ^24^ where: core gene families are present in 100% of isolates (N=22); softcore gene families are present in less than 100% but more than or equal to 95% of isolates (N=21); shell gene families are present in 5-95% of isolates (21>N≥2); cloud gene families are present in less than 5% of isolates (N=1). The putative function of each gene family was determined using by annotating the genes in each family using InterProScan, v5.53-87.0 ^62^ and the Pfam ^63^, PANTHER ^64^, and TIGRFAM ^65^ databases.

The patterns of core and accessory (softcore, shell, and cloud) biosynthetic gene cluster families were also examined. Biosynthetic gene cluster boundaries were predicted using antiSMASH, v4.1.0 ^66^. Two biosynthetic gene clusters were determined to be putatively orthologous if at least 50% of the genes they encode are reciprocally orthologous following a previously established protocol ^67^. The putative function of each biosynthetic gene cluster was determined by cross-referencing predicted clusters with known clusters in the MIBiG database ^68^.

### Phenotyping among infection-relevant traits

To determine growth on solid media, a drop containing 1 x 10^4^ spores was inoculated at the center of a petri dish containing solid minimal medium [1% (w/v) glucose, original high nitrate salts, trace elements, pH 6.5]. The plates were incubated at 37°C for 5 days. Trace elements, vitamins, and nitrate salts compositions were as described previously ^69^.

Radial growth was used to compare how the different strains respond to cell wall stressors (congo red and calco-fluor white-CFW), temperature (30, 37 e 44°C) and oxidative stress (H_2_O_2_ and menadione). Strains were grown in solid GMM, inoculated with 1X10^5^, and incubated for 5 days at 37°C before colony diameter was measured. To induce cell wall stress, 40 μg/mL of Congo red (Sigma-Aldrich) or CFW (Sigma-Aldrich) was added to medium. For oxidative stress assay, plates supplemented with 0.1 mM menadione (Sigma-Aldrich) and 10mM H_2_O_2_ (Merck S. A) were used. Radial growth for the aforementioned stresses was expressed as ratios, dividing colony radial diameter (cm) of growth in the stress condition by colony radial diameter in the control (no stress) condition. Finally, the radial growth of the strains were quantified at 30 and 44°C as well.

Antifungal susceptibility testing for voriconazole (Sigma-Aldrich), amphotericin B (Sigma-Aldrich) and caspofungin (Sigma-Aldrich) was performed by determining the minimal inhibitory concentration (MIC) or minimal effective concentration (MEC) (only caspofungin) according to the protocol established by the Clinical and Laboratory Standards Institute ^70^.

The diameters of 100 spores for each isolate were measured under a Carl Zeiss (Jena, Germany) AxioObserver.Z1 fluorescent microscope equipped with a 100-W HBO mercury lamp using the 100x magnification oil immersion objective and the AxioVision, software v.3.1.

For the crossing experiments, the lineages were inoculated minimal medium and incubated for two days at 37°C. After that, for sex cycle induction, the plates were sealed with tape to reduce oxygen tension. After incubation for 14 days at 30°C and 37°C, the cleistothecia were isolated using a magnifier and cleaned in a petri dish containing 4% (p/v) of agar. Cleistothecia were broken into microtubes containing water in order to release the ascospores, which were inoculated in minimal medium and incubated at 37°C to check their viability.

To visualize phenotypic variation between strains of *A. latus*, A*. spinulosporus*, *A. quadrilineatus*, and *A. nidulans*, a principal component analysis (PCA) using the factoextra, v2.8 (https://cran.r-project.org/web/packages/factoextra), and FactoMineR, v1.0.7 (https://cran.r-project.org/web/packages/FactoMineR), packages in R, v4.3.0, was utilized. More specifically, the dataset included measures of fungal growth in response to cell wall (Calcofluor White and Congo red at 40 μg/mL), oxidative (menadione, at 0.07 mM and 0.1 mM concentrations, and H_2_O_2_ for three days), and temperature (44°C, 37°C, and 30°C) stressors, in addition to various antifungal medications (voriconazole, amphotericin B, and caspofungin). Other physical characteristics in the growth of each isolate, such as asexual spore (conidial) size and the presence or absence of a viable cleistothecium, were also included for a total of 14 variables used in this PCA. Data were scaled to unit variance before the analysis (scale.unit = TRUE), and 5 dimensions were kept in the final results (ncp = 5).

### Survival Curves

*Galleria mellonella* larvae were used to investigate the virulence of the different strains. The larvae were cultivated and prepared as described previously ^20^. The larvae used for the infection were in the last stage of development (sixth week). All selected larvae weighed ∼300 mg and were restricted to food for 24 h before the experiment. The fresh conidia of each specie were counted using a hemocytometer and adjusted to the final concentration of 2 × 108 conidia/mL. Five microliter of each inoculum was injected with a Hamilton syringe (7000.5 KH) through the last left ear (n=10/group), resulting in 1 × 106 conidia. The control group was inoculated with phosphate buffered saline (PBS). After infection, the larvae were kept with food restrictions, at 37°C in Petri dishes in the dark and scored daily. The larvae were considered dead due to lack of movement in response to touch. The viability of the inoculum administered was determined by serial dilution of the conidia in YAG medium and incubating the plates at 37°C for 72 h. The experiment was repeated twice. We separated and assembled the groups with the larvae (n = 10) in Petri dishes. The groups are composed of larvae that are approximately 300 mg in weight and 2 cm long. Moth sex was not accounted for due to the impossibility of visually determining sex at the sixth week of larval development.

### Fermentation, Extraction, and Isolation for Chemotyping

To identify the secondary metabolites biosynthesized by *A. nidulans*, *A. latus*, *A. quadrilineatus*, and *A. spinulosporus*, these species were grown as large-scale fermentations to isolate and characterize the secondary metabolites, similar to methods previously described ^71^. To inoculate oatmeal cereal media (Old Fashioned Breakfast Quaker oats), agar plugs from fungal stains grown on potato dextrose agar; Difco were excised from the edge of the Petri dish culture and transferred to separate liquid seed medium that contained 10 ml YESD broth (2% soy peptone, 2% dextrose, and 1% yeast extract; 5 g of yeast extract, 10 g of soy peptone, and 10 g of D-glucose in 500 ml of deionized H2O) and allowed to grow at 23 °C with agitation at 100 rpm for 3 days. The YESD seed cultures of the fungi were subsequently used to inoculate solid-state oatmeal fermentation cultures, which were either grown at room temperature (∼23 °C under 12 hr light/dark cycles for 16 days), 30 °C for 12 days, or 37 °C; all growth at the latter two temperatures was carried out in an incubator (VWR International). The oatmeal cultures were prepared in 250 ml Erlenmeyer flasks that contained 10 g of autoclaved oatmeal (10 g of oatmeal with 17 ml of deionized H2O and sterilized for 15–20 min at 121 °C). For all fungal strains, six flasks of oatmeal cultures were grown at 37 °C and 30 °C, except for *A. quadrilineatus* which wasn’t grown at 37 ° C. Only one flask of each strain was grown for the growths at room temperature.

The cultures were extracted by adding 60 ml of 1:1 mixture of MeOH-CHCl_3_, chopping thoroughly with a spatula, and shaking overnight (∼16 hr) at ∼100 rpm at room temperature. The culture was filtered *in vacuo*, and 90 ml CHCl_3_ and 150 ml H_2_O were added to the filtrate. The sample was transferred to a separatory funnel and the organic layer was drawn off and evaporated to dryness *in vacuo*. The dried organic layer was reconstituted in 100 ml of (1:1) MeOH–CH_3_CN and 100 ml of hexanes, transferred to a separatory funnel, and shaken vigorously. The defatted organic layer (MeOH–CH_3_CN) was evaporated to dryness *in vacuo*.

### Sterigmatocystin Calibration Curve

High-resolution electrospray ionization mass spectrometry (HRESIMS) experiments utilized a Thermo LTQ Orbitrap mass spectrometer (Thermo Fisher Scientific) that was equipped with an electrospray ionization source. This was coupled to an Acquity ultra-high-performance liquid chromatography (UHPLC) system (Waters Corp.), using a flow rate of 0.3 ml/min and a BEH C_18_ column (2.1 mm x 50 mm, 1.7 μm) that was operated at 40°C. The mobile phase consisted of CH_3_CN–H_2_O (Fischer Optima LC-MS grade; both acidified with 0.1% formic acid). The gradient started at 15% CH_3_CN and increased linearly to 100% CH_3_CN over 8 min, where it was held for 1.5 min before returning to starting conditions to re-equilibrate.

Extracts were analyzed with four biological replicates, and each of these in technical triplicates, all in the positive ion mode with a resolving power of 35,000. The samples were each prepared at a concentration of 0.2 mg/ml and were dissolved in MeOH and injected with a volume of 3 μl. To eliminate the possibility for sample carryover, a blank (MeOH) was injected between every sample. The sterigmatocystin calibration curve was analyzed in triplicate and ran at concentrations of 102.4, 51.2, 25.6, 6.4, 3.2, 1.6, 0.8 μg/mL. To ascertain the absolute concentration of sterigmatocystin in these samples, batch process methods were run using Thermo Xcalibur (Thermo Fisher Scientific). We used a mass range of 325.0687-325.0719 Da for sterigmatocystin at a retention time of 5.00 min with a 7.00 second window. The calibration curve was analyzed quadratically with a weight index of 1/X since these compounds did not ionize linearly at this large concentration range. The equation used to calculate the absolute concertation of sterigmatocystin in these samples was, Y = 1.99463e+007+5.0315e+007*X-212053*X^2^ with a R^2^ of 0.9987. The absolute concentration of sterigmatocystin per flask of fungal growth was calculated by converting the concentration in the sample injection to the amount in the total extract, followed by averaging the biological replicates.

### Chemometric Analysis

Untargeted UPLC-MS datasets for each sample were individually aligned, filtered, and analyzed using MZmine 2.20 software (https://sourceforge.net/projects/mzmine/) ^72^. Peak detection was achieved using the following parameters: noise level (absolute value), 5×10^5^; minimum peak duration, 0.05 min; m/z variation tolerance, 0.05; and m/z intensity variation, 20%. Peak list filtering and retention time alignment algorithms were used to refine peak detection. The join algorithm integrated all sample profiles into a data matrix using the following parameters: m/z and retention time balance set at 10.0 each, m/z tolerance set at 0.001, and RT tolerance set at 0.5 mins. The resulting data matrix was exported to Excel (Microsoft) for analysis as a set of m/z – retention time pairs with individual peak areas detected in triplicate analyses. Samples that did not possess detectable quantities of a given marker ion were assigned a peak area of zero to maintain the same number of variables for all sample sets. Ions that did not elute between 2 and 8 minutes and/or had an m/z ratio less than 200 or greater than 800 Da were removed from analysis.

Relative standard deviation was used to understand the quantity of variance between the technical replicate injections, which may differ slightly based on instrument variance. A cutoff of 40% was used at any given m/z – retention time pair across the technical replicate injections of one biological replicate, and if the variance was greater than the cutoff, it was assigned a peak area of zero. The data underwent standard scaling to normalize the data. Final chemometric analysis was performed with Python. The network graphs were generated using the standard scaled data with Pandas.

### Flow cytometry analysis for determination of spores DNA content

Asexual spores were collected, centrifuged (13,000 rpm for 3 min), and washed with sterile PBS. Spores were stained following the protocol previously described ^73^. Briefly, after harvesting, spores were fixed overnight with 70% ethanol (v/v) at 4°C, washed and resuspended in 850 μl of sodium citrate buffer (50 mM; pH = 7.5) and dissociated by sonication using four ultrasound pulses at 40W for 2 seconds with an interval of 1 to 2 seconds between pulses. Spores were treated with RNase A (0.50 mg/mL; Invitrogen, USA; 1 hour; 50°C) and then with proteinase K (1 mg/mL; Sigma-Aldrich, St. Louis, Missouri, USA; 2 hours; 50°C). Spore DNA was stained with SYBR Green 10,000x (Invitrogen, USA; 0.2% (v/v)), overnight at 4°C. Prior to analysis, spores were treated with Triton® X-100 (Sigma-Aldrich; 0.25% (v/v)). Samples were acquired in an LSRII flow cytometer (Becton Dickinson, NJ, USA) with a 488-nanometer excitation laser with Diva as the acquisition software. A minimum of 30,000 spores per sample were analyzed with an acquisition protocol defined to measure forward scatter and side scatter on a logarithmic scale and green fluorescence on a linear scale. Results were analyzed with the FlowJo software (BD Biosciences, USA).

### RNA-sequencing extraction and analysis

The RNA extraction was performed according to dos Reis et al., 2021 with modifications ^74^. Briefly, 10^6^ spores/mL of each *Aspergillus* strain was inoculated in 50 mL of liquid minimal medium and grown at 37°C for 16 h under agitation. Further, the mycelia were filtered through miracloth, freeze dried, and disrupted by grinding in liquid nitrogen. Total RNA was extracted using TRIzol (Invitrogen), treated with RQ1 RNase-free DNase I (Promega) and purified using the RNAeasy kit (Qiagen) according to the manufacturer’s instructions. RNA samples were quantified using a NanoDrop and its RNA quality was verified using an Agilent Bioanalyzer 2100 (Agilent Technologies). RNAs selected for further analysis had a minimum RNA integrity number (RIN) value of 8.0. For RNA-sequencing, the cDNA libraries were constructed using the TruSeq Total RNA with Ribo Zero (Illumina, San Diego, CA, USA). From 0.1–1 μg of total RNA, the ribosomal RNA was depleted, and the remaining RNA was purified, fragmented, and prepared for complementary DNA (cDNA) synthesis, according to the manufacturer’s instructions. The libraries were validated following the Library Quantitative PCR (qPCR) Quantification Guide (Illumina). Following, the RNA-seq was carried out by paired-end sequencing on the Illumina NextSeq 500 Sequencing System using NextSeq High Output (2 x 150) kit, according to the manufacturer’s instructions. Using the same approach, RNA was also extracted from spores grown at 30°C.

RNA was sequenced at Vanderbilt Technologies for Advanced Genomics (VANTAGE) on an Illumina NovaSeq machine (paired-end, 150bp length for each read). All samples passed quality control checks and had acceptable RNA Integrity Numbers. Reads were trimmed with Trimmomatic, v0.39 ^75^, using the following options “TruSeq2-PE.fa:2:30:10 LEADING:3 TRAILING:3 SLIDINGWINDOW:4:15 MINLEN:36”. Filtered reads for each condition and replicate were then mapped to the hybrid genome assembly and annotation file using HISAT2, v2.1.0 ^76^, with default parameters. The resulting SAM files from HISAT2 were then sorted and compressed to BAM files with SAMtools, v1.9 ^77^. Gene abundance for each condition and replicate were estimated by first filtering the alignment files for only properly paired reads using SAMtools (samtools view -b -f 2 <input.bam> < input.proper.pairs.only.bam>). Per-sample gene abundances were then quantified in terms of transcripts per million (TPM) using TPMcalculator, v0.0.3 ^78^. To identify genes that appear differentially expressed between the two treatment conditions, we extracted per-sample unique read counts from each alignment file using from HTseq ^79^ and then passed these counts to DESeq2, v1.34 ^80^, to estimate differential expression between conditions. Genes were considered to be significantly differentially expressed if their FDR-corrected (Benjamini and Hochberg method) p-value (padj) was lower than 0.05.

## Data Availability

The genome sequencing and transcriptomic data have been submitted to the NCBI (BioProject accession PRJNA987017). Accessions for short-read and long-read genome sequences are SRR25010807-SRR25010890. Accessions for transcriptomic data are SRR25016725-SRR25016730. Key results and intermediate files—genome assemblies, annotations, orthology inference, and biosynthetic gene cluster boundaries, for example—have been uploaded to figshare (doi: 10.6084/m9.figshare.23589570).

## Acknowledgements

We thank the Rokas lab and Dr. Judith Berman for helpful discussion and feedback. J.L.S. and A.R. were funded by the Howard Hughes Medical Institute through the James H. Gilliam Fellowships for Advanced Study program. Research in A.R.’s lab is supported by grants from the National Science Foundation (DEB-2110404), the National Institutes of Health/National Institute of Allergy and Infectious Diseases (R01 AI153356), and the Burroughs Wellcome Fund. Research in G.H.G’s lab is supported by the Fundação de Amparo à Pesquisa do Estado de São Paulo (FAPESP) grants numbers 2016/07870-9 and 2021/04977-5 (G.H.G.) and the Conselho Nacional de Desenvolvimento Científico e Tecnológico (CNPq) grant numbers 301058/2019-9 and 404735/2018-5 (G.H.G.), both from Brazil. Research in F.R.’s Lab was supported by the European Union’s Horizon 2020 research and innovation program under grant agreement no. 847507 (HDM-FUN).

