## Supplementary Figures and Tables for "Evolutionary origin, population diversity, and diagnostics for a cryptic hybrid pathogen"

**Extended Data for**

### The evolutionary origins and phenotypic impact of hybridization in a cryptic fungal pathogen

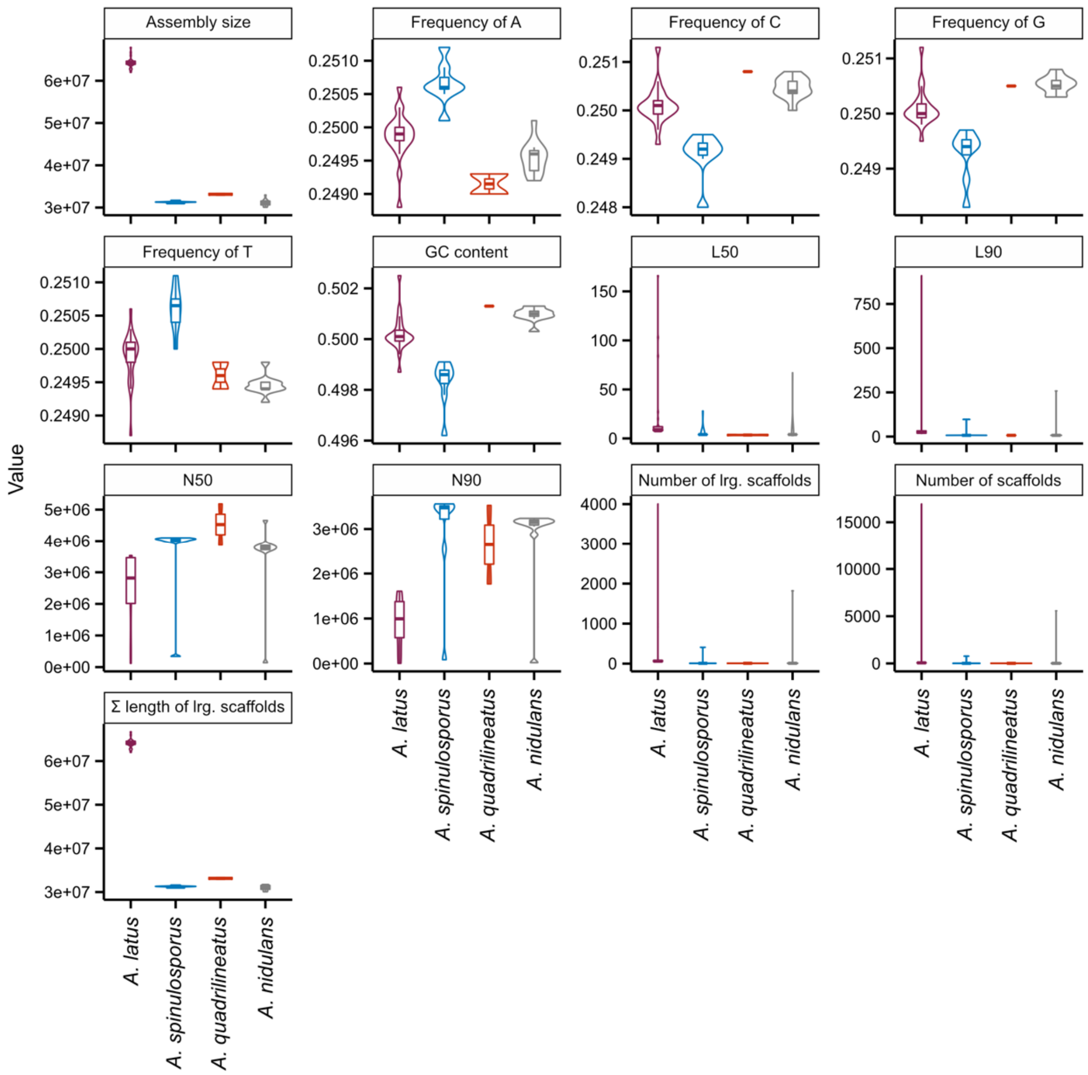


**Extended Data Fig. 1. Summary of genome assembly metrics among 41 clinical isolates.** Each panel depicts a different metric of genome assembly quality or feature. From top left to bottom right, each panel depicts assembly size, the frequency of A, C, G, and T nucleotides, GC content, L50, L90, N50, N90, the number of large scaffolds (>500 base pairs), the number of scaffolds, and the sum length of large scaffolds. The high L50 and L90 values as well as the number of scaffolds and low N50 and N90 values were observed in the genome assembled using only short-read data, MO46149, which was obtained from a previous study (Steenwyk *et al.* 2020).

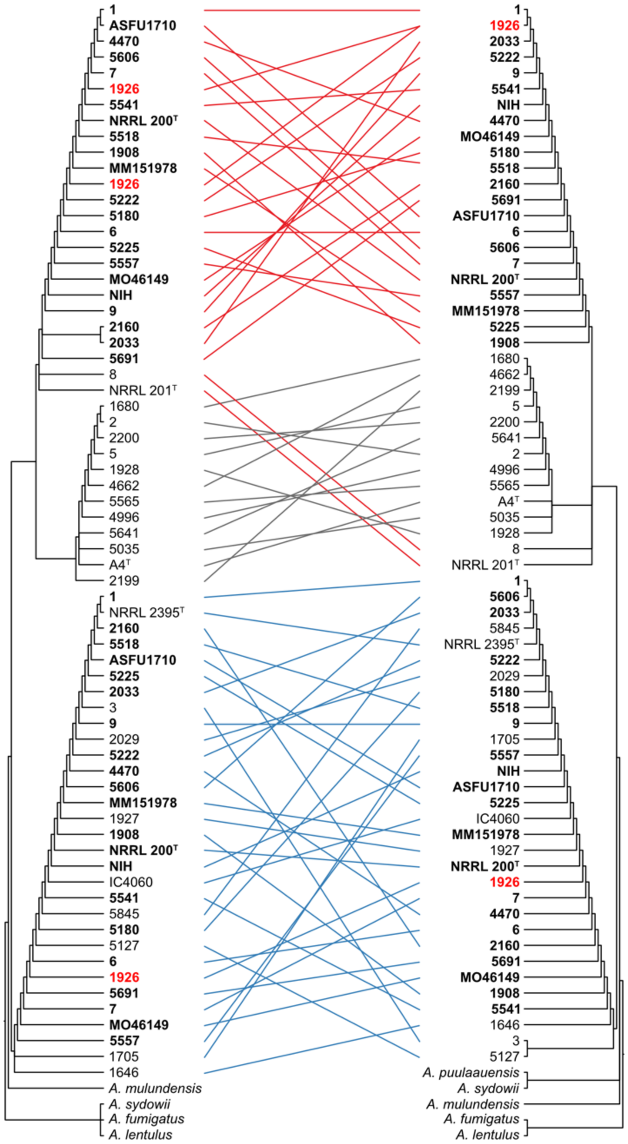


**Extended Data Fig. 2. Molecular phylogenetics of taxonomically informative loci.** Phylogenies of β-tubulin (left) and calmodulin sequences are depicted as a tanglegram wherein lines inbetween the phylogenies are drawn between the same two isolates. Isolates with two copies of each locus are shown in bold. Note, isolate 1926 had three copies of β-tubulin and is flagged in red. Phylogenies are depicted without branch length information. Branch lengths were often zero (or close to zero) resulting in ladder-like topology. Lines in blue indicate sequences monophyletic with *A. spinulosporus* NRRL 2395$^T$, grey indicates sequences monophyletic with *A. nidulans* A4, red indicates sequences that monophyletic or closely related to *A. quadrilienatus* NRRL 201$^T$.

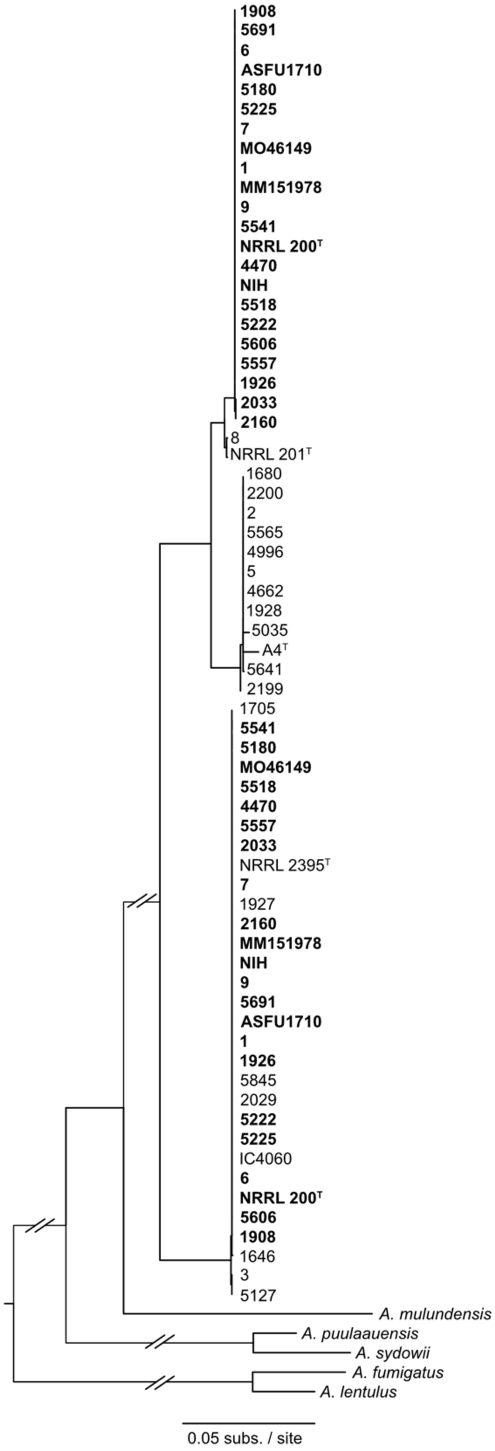


**Extended Data Fig. 3. Molecular phylogenetics of concatenated taxonomically informative loci.** The phylogeny inferred from concatenating the β-tubulin and calmodulin sequence is depicted here. Branch lengths represent substitutions per site. Isolates that have two copies of each locus are shown in bold.

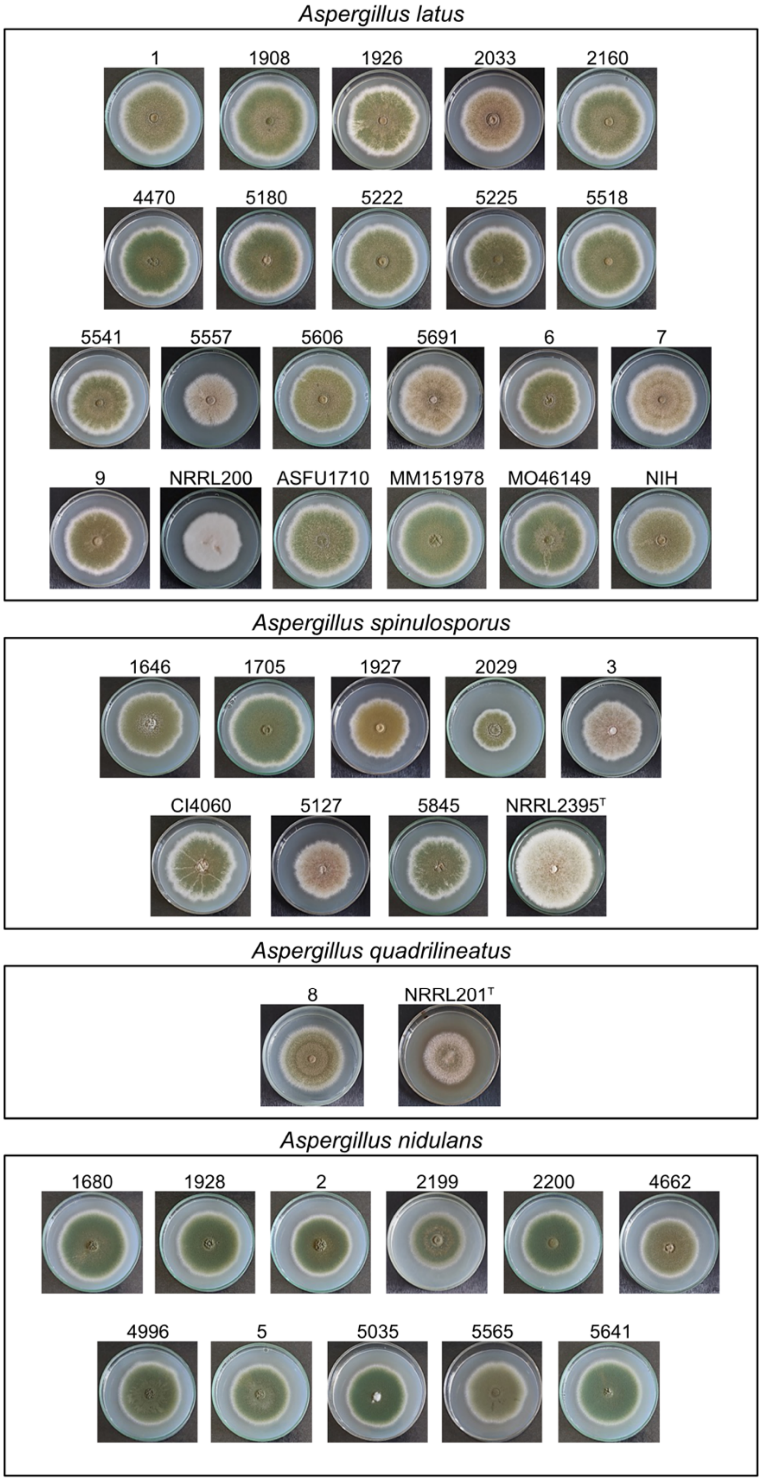


**Extended Data Fig. 4. Isolates used in the present study are difficult, if not impossible, to visually bin into species.**

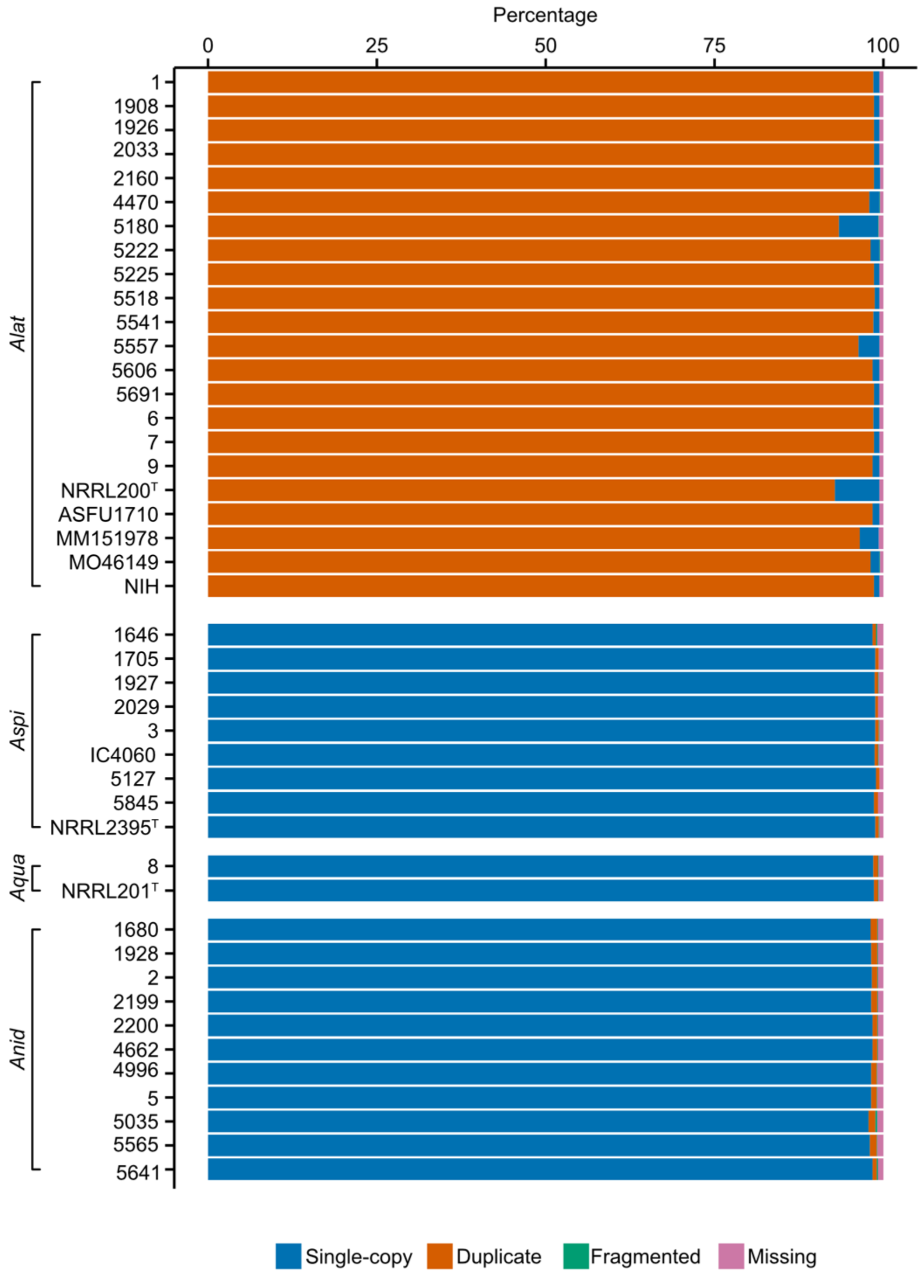


**Extended Data Fig. 5. *Aspergillus latus* genomes encode many duplicated BUSCO genes.** Isolates of *Aspergillus latus* have an average of 97.79 ± 1.65% duplicated BUSCO genes. The y-axis depicts various isolates examined in the present study organized by species. The x-axis depicts the percentage of BUSCO genes that single-copy (blue), duplicated (orange), fragmented (green), and missing (pink). *Alat* corresponds to *A. latus*; *Aspi* corresponds to *Aspergillus spinulosporus*; *Aqua* corresponds to *Aspergillus quadrilineatus*; and *Anid* corresponds to *Aspergillus nidulans*. The Eurotiales database of BUSCO genes was used (Creation date: 2020-08-05, number of species: 60, number of BUSCOs: 4191).

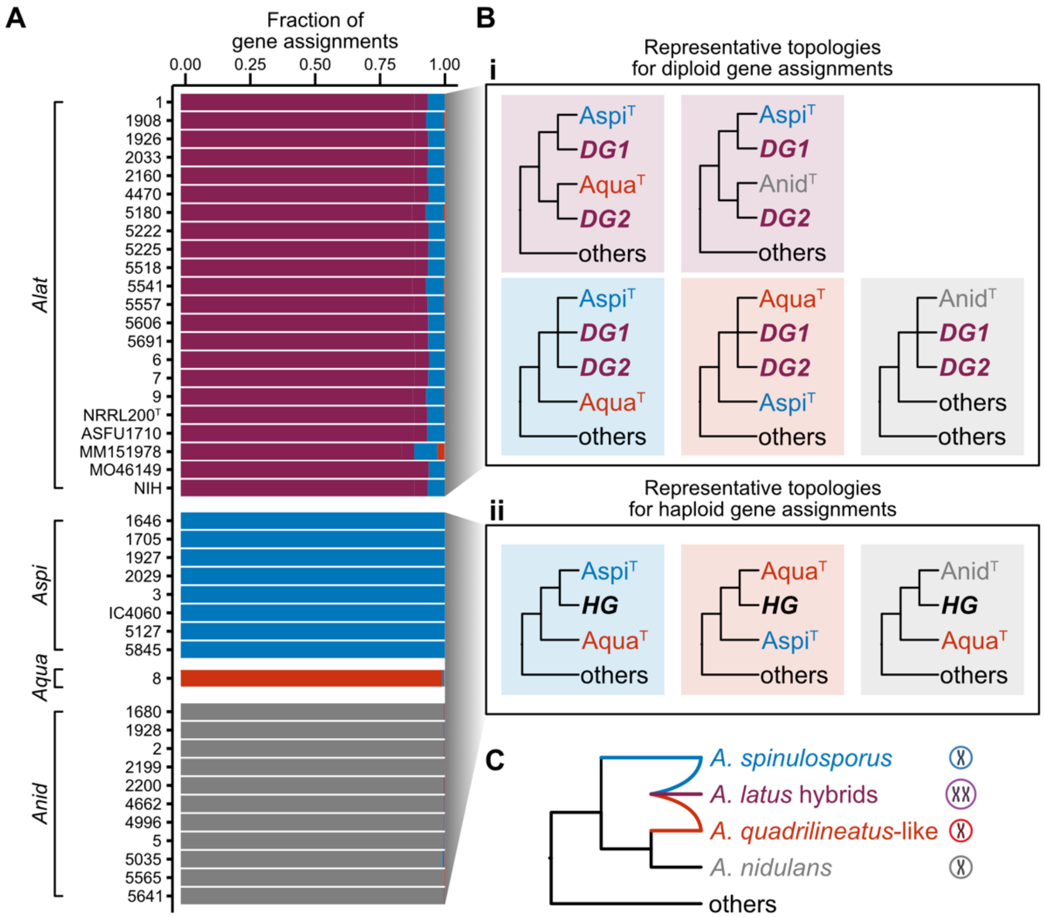


**Extended Data Fig. 6. Phylogenetic determination of gene parent-of-origin confirms *Aspergillus latus* arose via hybridization.** (A) Isolates are shown along the y-axis. The x-axis depicts the fraction of gene assignments that reflected different single-gene phylogenetic topologies. (B) Cartoon representations of different single-gene phylogenetic topologies are shown here. (i) For diploid genomes, purple topologies revealed one gene (depicted as DG1) was closely related to *Aspergillus spinulosporus* whereas the other gene copy (depicted as DG2) was more closely related to *Aspergillus quadrilineatus* or *Aspergillus nidulans*. Some topologies were observed in which both diploid genes were more closely related to *A. spinulosporus*, an observation that may be driven by lack of taxon sampling of the species most closely related to the second sequence. (ii) A similar approach to gene assignments was taken for haploid genomes, but only one gene (depicted as HG) was examined. (C) These findings together with other observations support a hybridization event between *A. spinulosporus* and a close relative of *A. quadrilineatus* (or *A. quadrilineatus*-like) gave rise to *A. latus*.

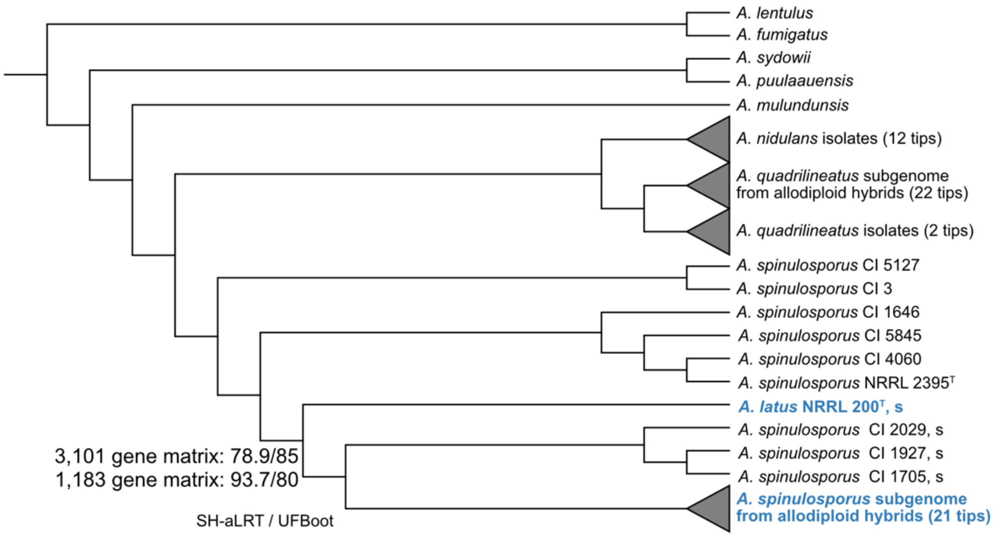


**Extended Data Fig. 7. Two hybridization events are not well supported.** Examination of the bipartition in which *A. latus* NRRL 200$^{T}$ diverged from the 21 other *A. spinulosporus* subgenomes of the hybrid species (shown in blue) and three clinical isolates of *A. spinulosporus* revealed non-optimal support using a 3,101 gene matrix. Subsampling the complete dataset for genes with taxon occupancy equal to or greater than 38 among hybrid subgenomes revealed support for two hybridization events was still non-optimal, albeit more robust. Support is only depicted for the bipartition of interest. Support was evaluated using two measures: SH-aLRT and UFBoot.

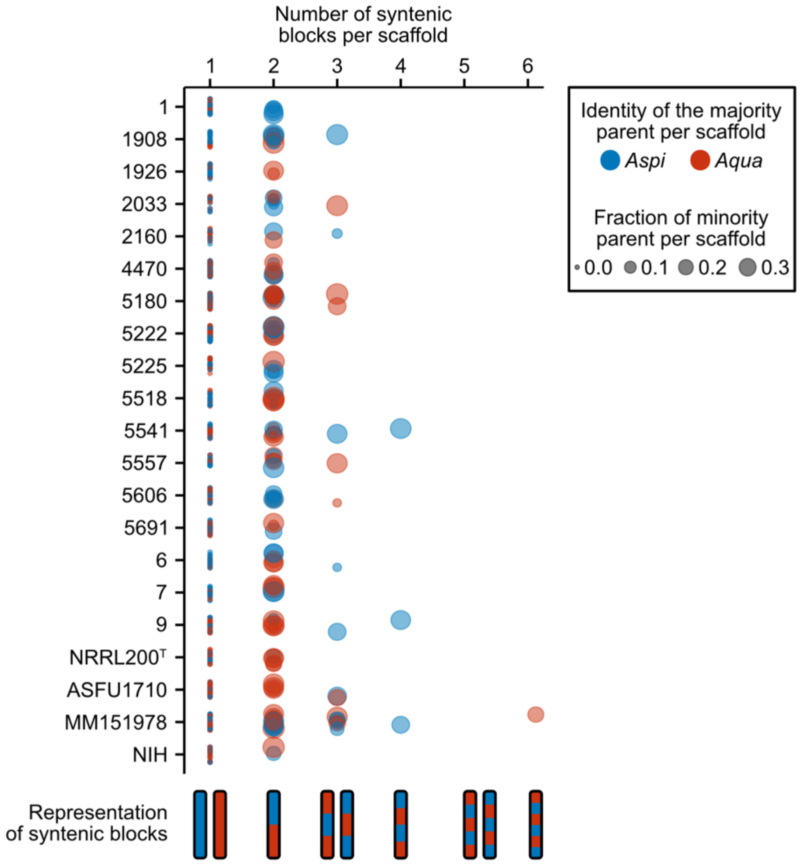


**Extended Data Fig. 8. The number of syntenic blocks per scaffold.** Each data point represents one scaffold. Syntenic blocks are defined as having 15 or more consecutive genes assigned to one parent in a hybrid genome. Scaffolds may have one syntenic block where all genes on the scaffold were assigned to one parent. Scaffolds with putative evidence of recombination have two or more syntenic blocks. Up to six syntenic blocks were observed in a scaffold. For each scaffold, the majority parent is denoted by the data point color. The datapoint size reflects that fraction of genes assigned to the minority parental scaffold; thus, large data points have nearly even numbers of genes from each parent whereas small data points have few genes from the parent with less representation on the scaffold.

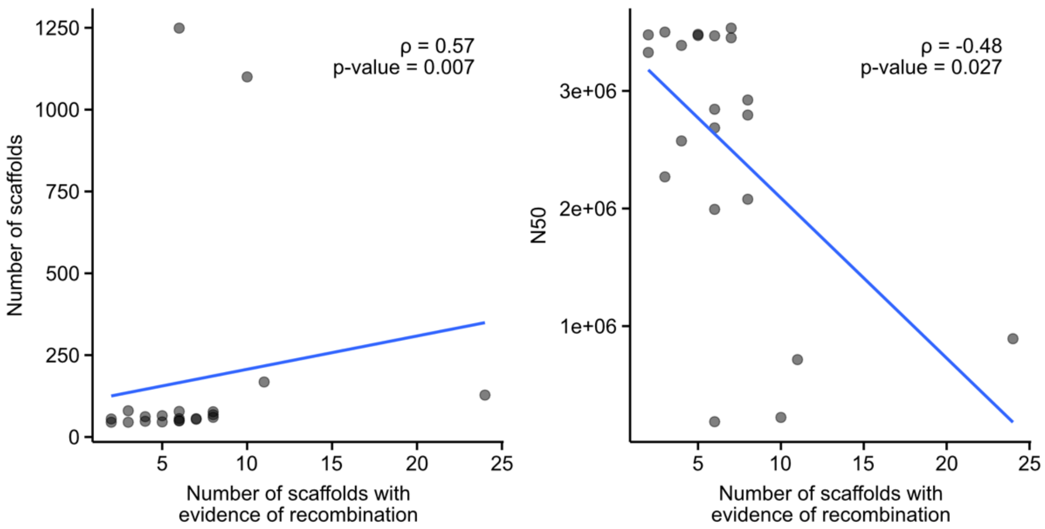


**Extended Data Fig. 9. The number of putatively recombinant scaffolds increases as genome assembly quality decreases.** (Left) A significant positive correlation between the number of scaffolds with evidence of recombination and the number of scaffolds in a genome assembly. (Right) A significant negative correlation was observed between the number of scaffolds with evidence of recombination and genome assembly N50, a measure of contiguity.

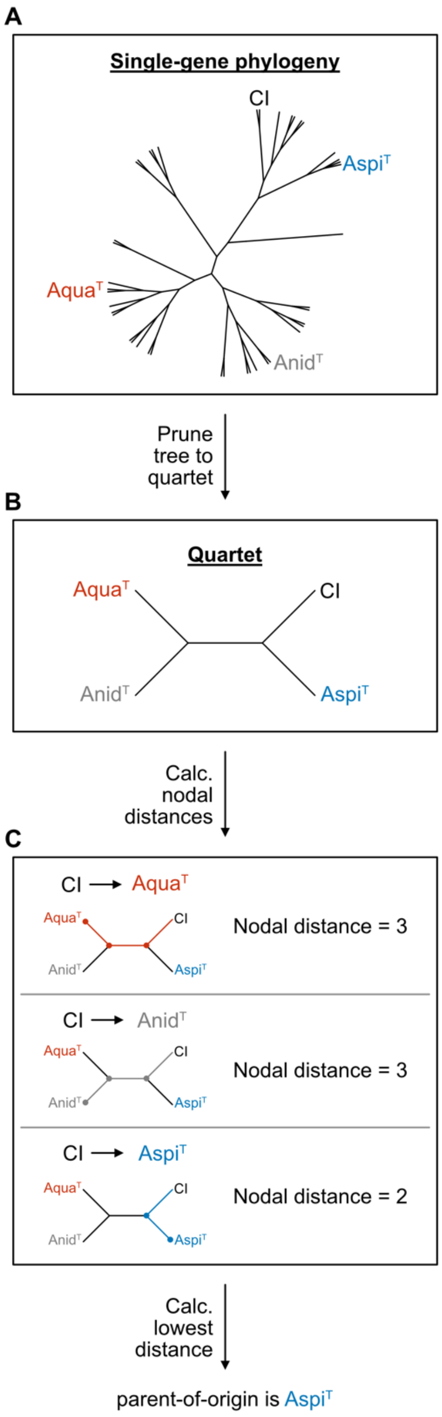


**Extended Data Fig. 10. Tree-based assignment of genes to parent-of-origin.** (A) Large phylogenies that include single-copy sequences from *A. spinulosporus* NRRL2395$^T$, *A. quadrilineatus* NRRL201$^T$, and *A. nidulans* A4 as well as duplicate-copy sequences from hybrid strains and single-copy sequences from haploid strains represented as CI (clinical isolate) were inferred. The resulting phylogenies were pruned (or decomposed) into a quartet with *A. spinulosporus* NRRL2395$^T$, *A. quadrilineatus* NRRL201$^T$, *A.*

*nidulans* A4, and one sequence from a CI. (C) Nodal distances were calculated between the CI sequence and *A. spinulosporus* NRRL2395$^T$, *A. quadrilineatus* NRRL201$^T$, and *A. nidulans* A4. Parent-of-origin was determined by the shortest nodal distance. In this case, the parent-of-origin is *A. spinulosporus* NRRL2395$^T$.

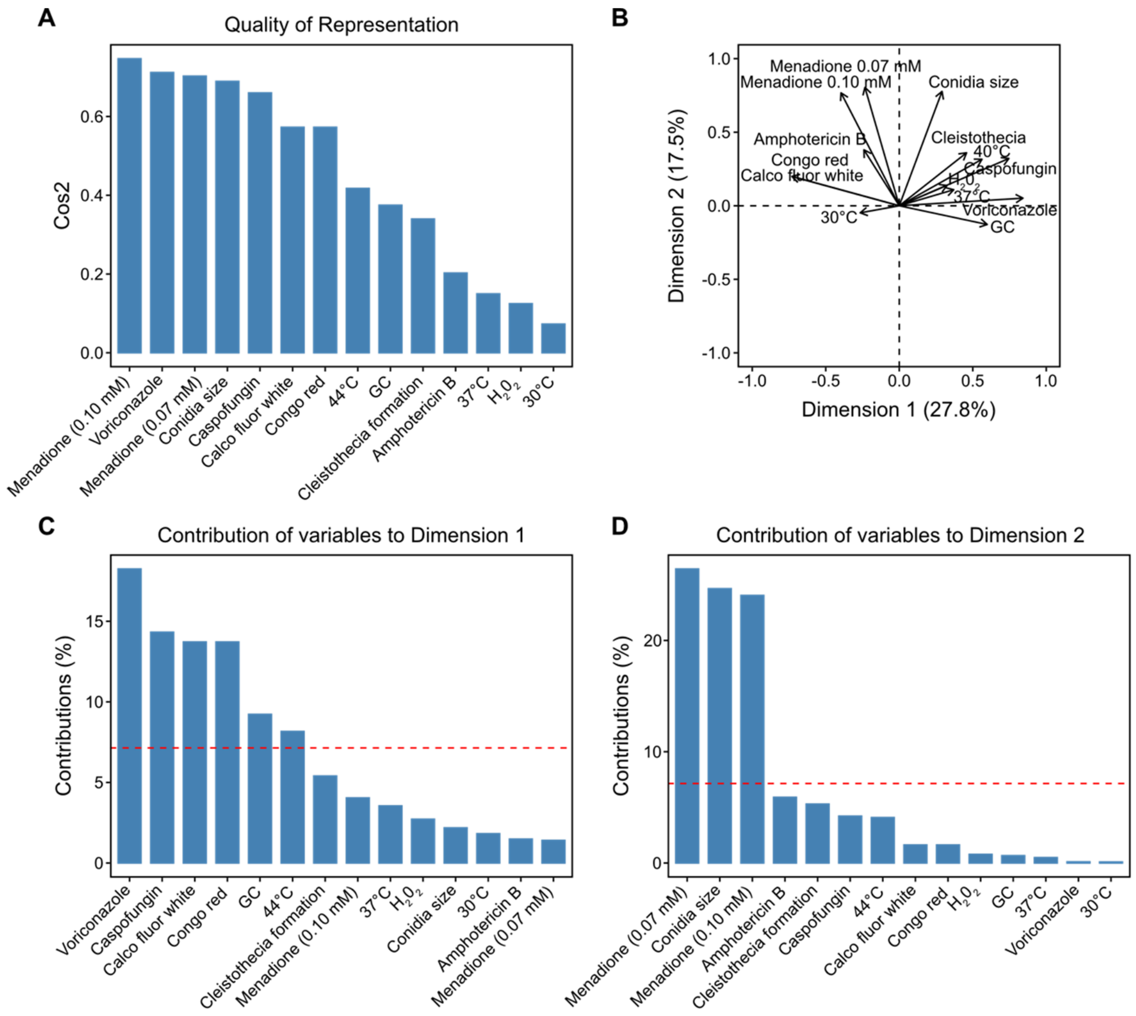


**Extended Data Fig. 11. Quality, correlations, and contributions of phenotypes to principal component space.** (A) Quality of variable representation measured using Cos2 (square cosine) along all principal components. (B) Correlations among phenotypes, which are depicted as arrows. Arrows pointing in the same direction are correlated (e.g., growth in Menadione at concentrations of 0.10 mM and 0.07 mM) whereas arrows pointing in the opposite direction are anti-correlated (e.g., the minimum inhibitory concentration of voriconazole and growth at 30°C). The distance between the phenotype arrows and the origin of the circle reflects Cos2 values. (C) Examination of contributions of variables to dimension one of the principal component space reveals Voriconazole contributes the most. (D) Examination of the contribution of variables to dimension two of the principal component space reveals that growth in Menadione (0.07 mM) contributes the most.

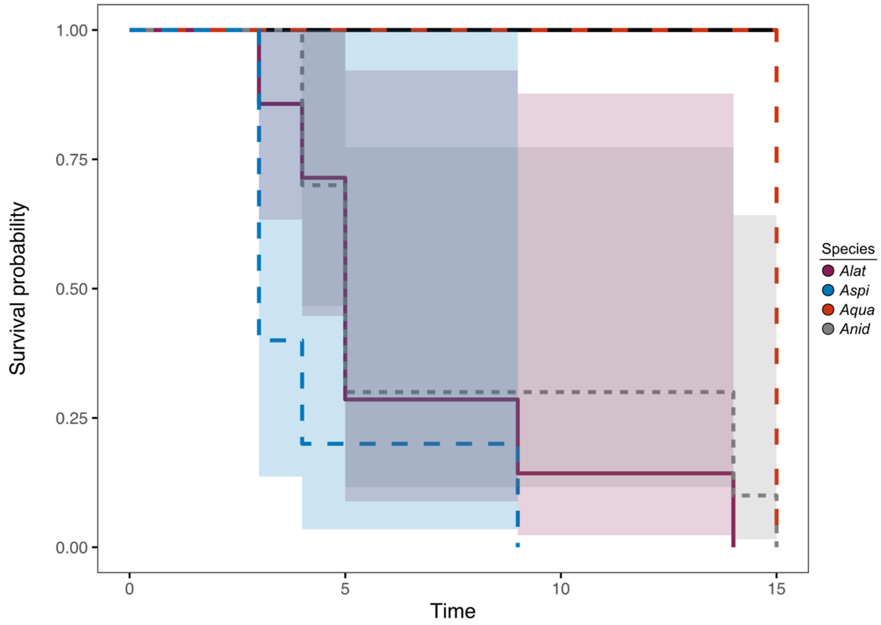


**Extended Data Fig. 12. Survival curves of an invertebrate moth model of disease.** Survival probabilities of invertebrate moths are depicted for each species wherein multiple strains were tested. Survival curves among the various species were significantly different (p = 0.04; Log-Rank test). More specifically, *A. spinulosporus* is the most pathogenic followed by *A. latus* and *A. nidulans*, which had similar survival curves. Lastly, *A. quadrilineatus,* was the least virulent. Note, for *A. quadrilineatus*, only one strain as tested and therefore no confidence intervals have been included. A PBS control is depicted as a black line.

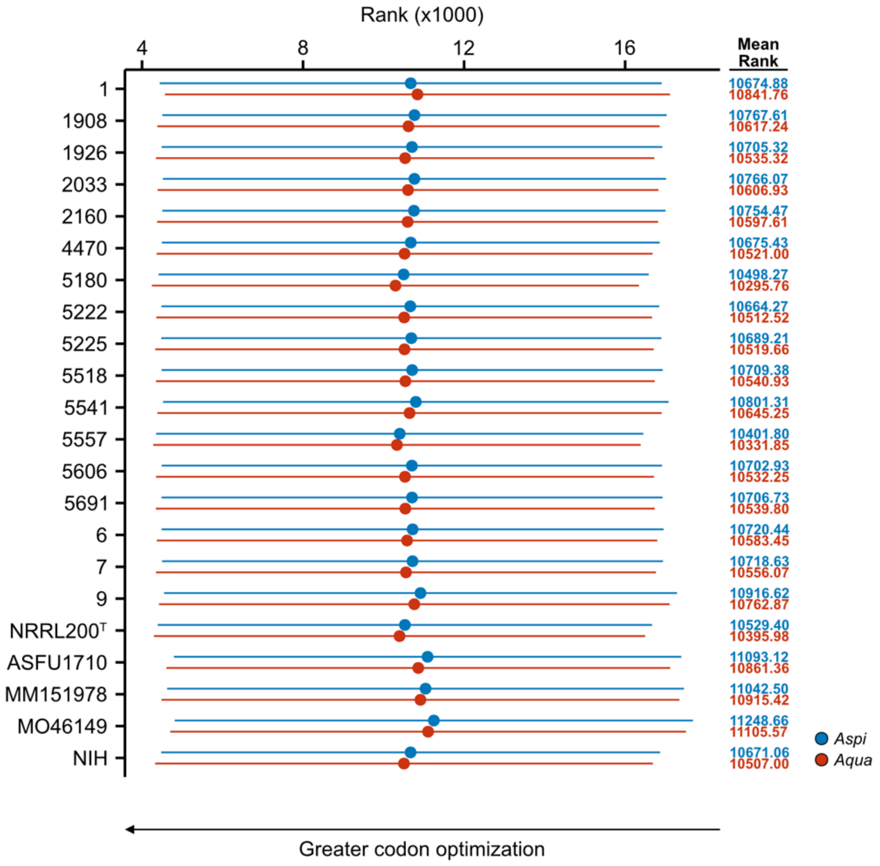


**Extended Data Fig. 13. Codon optimization suggests the *Aspergillus quadrilineatus*-like parental genome is more optimized.** Codon optimization was estimated using measures of mean gene-wise relative synonymous codon usage (gRSCU). Genes were ranked from most codon optimized to least codon optimized using the gRSCU values. Genes were assigned to each parent using a reciprocal best BLAST hit approach. The average rank plus/minus one standard deviation is plotted. Across all isolates the *A. quadrilineatus*-like subgenome is more codon optimized than the *Aspergillus spinulosporus* subgenome. Examination of the differences in codon optimization for each parent among allodiploid hybrid genomes revealed a significant different between the optimization of the parental subgenomes and isolates ($p < 0.001$ for both comparisons, Multi-factor ANOVA). No interaction was observed between parental subgenome and isolates ($p = 0.81$, Multi-factor ANOVA), thus, an additive model was used to examine if there are differences in the optimization of parental subgenome. Significant differences in parental subgroup optimization was observed ($p < 0.001$, Tukey Honest Significant Differences Test).

#### Supplementary Tables

| Description | Value |
|---|---|
| **LogL single hybridization event** | -26,608,690.95 |
| **LogL two hybridization events** | -26,608,862.42 |
| **LogL difference** | 171.47 |
| **Bootstrap proportion using RELL method** | 0.059 |
| **P-value of one-sided Kishino-Hasegawa test** | 0.055 |
| **P-value of Shimodaira-Hasegawa test** | 0.055 |
| **P-value of a weighted Kishino-Hasegawa test** | 0.055 |
| **P-value of a weighted Shimodaira-Hasegawa test** | 0.055 |
| **Expected likelihood weight** | 0.059 |
| **P-value of approximately unbiased test** | 0.071 |

**Supplementary Table 1. Topology test results using the 3,101-gene data matrix.** Topology testing was used to determine the number of hybridization events. The null hypothesis is that two hybridization events and a single hybridization event are equally likely. Multiple testing strategies were used and each failed to reject the null hypothesis. LogL: log likelihood.

| Description | Value |
|---|---|
| **LogL single hybridization event** | -10,250,368.53 |
| **LogL two hybridization events** | -10,250,526.52 |
| **LogL difference** | 157.98 |
| **Bootstrap proportion using RELL method** | 0.057 |
| **P-value of one-sided Kishino-Hasegawa test** | 0.055 |
| **P-value of Shimodaira-Hasegawa test** | 0.055 |
| **P-value of a weighted Kishino-Hasegawa test** | 0.055 |
| **P-value of a weighted Shimodaira-Hasegawa test** | 0.055 |
| **Expected likelihood weight** | 0.057 |
| **P-value of approximately unbiased test** | 0.060 |

**Supplementary Table 2. Topology test results using the 1,183 -gene data matrix.** An alternative and more complete data matrix was also used to evaluate the number of hybridization events. The null hypothesis is that two hybridization events and a single hybridization event are equally likely. Multiple testing strategies were used and each failed to reject the null hypothesis. LogL: log likelihood.
